# Robust fluorescent labeling and tracking of endogenous non-repetitive genomic loci

**DOI:** 10.1101/2025.08.22.671818

**Authors:** Alex Raterink, Rajarshi P. Ghosh, Linfeng Yang, Quanming Shi, Minh Khue Nguyen, Isaac B. Hilton, Jan T. Liphardt, Anna-Karin Gustavsson

**Author notes:** These authors contributed equally (Alex Raterink, Rajarshi P. Ghosh). Corresponding authors: Rajarshi P. Ghosh, and Anna-Karin Gustavsson.

## Abstract

The spatial organization and dynamics of a genome are central to gene regulation. While a comprehensive understanding of chromatin organization in the human nucleus has been achieved using fixed-cell methods, measuring the dynamics of specific genomic regions over extended periods in individual living cells remains challenging. Here, we present a robust and fully genetically encoded system for fluorescent labeling and long-term tracking of any accessible non-repetitive genomic locus in live human cells using fluorogenic and replenishable nanobody array fusions of the *Staphylococcus aureus* dCas9, and compact polycistronic single guide (sg)RNAs. First, we characterize the selectivity and photostability of our probes, enabling genome-wide visualization of chromatin dynamics at locally repetitive elements. Next, through multiplexed expression of 8–10 sgRNAs from polycistronic cassettes, we demonstrate efficient and sustained labeling of non-repetitive loci, enabling high-fidelity tracking of gene-proximal regions at exceptional spatial and temporal resolution. Finally, by correlating chromatin mobility with transcriptional activity at multiple genes, we find that local chromatin dynamics at 20 Hz are gene-specific and not necessarily dependent on transcription. Our approach is versatile, minimally invasive, and scalable, enabling multiplexed imaging of regulatory element dynamics involved in gene control, with broad applicability across diverse biological systems and disease contexts.

## Introduction

Chromatin structure and dynamics are intimately linked to gene regulation, although the exact relationship at the single-cell level during interphase remains unclear^1^. Dysregulation of the dynamic organization of chromatin can lead to aberrant gene expression patterns and has been implicated in a range of diseases, including cancers, neurological disorders, and autoimmune conditions^2–4^. Studies of chromatin architecture in fixed cell populations have revealed that chromatin organization is highly dynamic, adapting in response to gene expression programs specific to cell type and changing progressively during cellular differentiation^5,6^. Although single-cell approaches have highlighted the heterogeneity of chromatin structural states across cell populations, they offer limited insights into how these structures evolve within individual cells^7,8^.

Live-cell fluorescence imaging methods have been instrumental in elucidating the relationship between rapid, nanoscale chromatin dynamics and transcriptional regulation^9^. Fluorescent tagging of specific genomic loci in live cells has been made possible by innovative strategies, including the insertion of operator arrays near target loci, which are recognized by fluorophore-conjugated binding proteins, as well as the use of programmable DNA-binding platforms such as catalytically inactive Cas9 (dCas9) and transcription activator-like effectors (TALEs)^10–13^. These strategies offer high spatial resolution and are readily compatible with multiplexed and time-lapse imaging modalities, facilitating dynamic studies of chromatin architecture in living cells. Fluorescent-repressor operator systems (FROS), such as TetO arrays, yield a high signal-to-background ratio (SBR), facilitating precise detection of labeled loci. However, they necessitate stable genomic integration through gene editing, which can perturb local chromatin structure, potentially altering gene expression or chromatin dynamics. Additionally, the requirement for customized array insertion at each target site limits scalability and throughput, making the simultaneous or sequential imaging of multiple genomic loci technically demanding and labor-intensive^12–21^. TALEs offer targeting flexibility, but engineering sequence-specific TALEs has become less common given the relative simplicity and cost-effectiveness of designing single guide (sg)RNAs in the dCas9 system^22,23^. However, dCas9 systems often suffer from low SBR and rapid photobleaching, and targeting is typically limited to locally repetitive sites in the genome^11,24–28^. CRISPR-Tag and TriTag combine the FROS and dCas9 approaches but suffer from the same inflexibility as FROS systems^29,30^. Several studies have successfully targeted non-repetitive genomic loci using dCas9 or dCas12-based imaging approaches^11,31–36^. However, many of these methods rely on deploying large numbers of sgRNAs per locus (36-73^11^, 36-48^37^, 12-18^33^, 20^38^, 48-60^39^, 67^35^, 288^40^, 96-375^36^), significantly increasing experimental complexity and labor. Alternatively, single-sgRNA strategies^34,41,42^ are more streamlined but are susceptible to off-target binding, which can produce false-positive signals unless dCas9 levels are precisely controlled and sgRNA sequences are carefully optimized^43–46^. Non-repetitive genomic loci have been labeled using as few as 4 sgRNAs containing 14× MS2 loops; however, this strategy has so far been limited to single time-point imaging of a single gene in fixed cells^31^. Another study^32^ achieved allele-specific labeling using just 2–3 sgRNAs with 3–6 aptamer loops, a striking contrast to other reports that struggled to image low or non-repetitive regions, even with 8 or more aptamers per sgRNA^25,28,47–49^. Despite these advances, aptamer-based methods often produce non-specific nuclear puncta, attributed to the accumulation of unbound or degraded sgRNAs^49,50^. A distinct approach using dual-FRET molecular beacons enabled labeling with just 3 sgRNAs^51^. However, the requirement for beacon electroporation shortly before imaging may disrupt cellular physiology, and the reliance on dual-color detection further complicates multiplexed imaging. These limitations underscore the need for a minimally invasive, low-copy sgRNA, and highly specific live-cell labeling strategy compatible with dynamic and multiplexed imaging of non-repetitive genomic loci. We previously developed dSpCas9-ArrayG/N, a system in which *Streptococcus pyogenes* dCas9 (dSpCas9) is fused to a nanobody array, enabling high-precision, three-dimensional (3D) tracking of a repetitive chromosomal locus over thousands of imaging frames with spatial resolution in the tens-of-nanometers range^52^. However, to date, this method has been applied only to the labeling and tracking of a single, locally repetitive genomic region, limiting its broader applicability.

In this study, we present a robust, fully genetically encoded and scalable platform for long-term, high-resolution tracking of non-repetitive genomic loci in living cells. Leveraging the highly target-specific *Staphylococcus aureus* dCas9 (dSaCas9) and polycistronic sgRNA expression, our system enables efficient labeling of endogenous, non-repetitive sites using only 8–10 sgRNAs transcribed from a single cassette. We first systematically characterize the binding kinetics of dSaCas9 and evaluate the photostability of our nanobody-based fluorescent amplifier (ArrayG/N), establishing the foundation for sustained, precise imaging. We then apply this platform to quantify locus-specific chromatin dynamics at multiple repetitive regions genome-wide and extend its utility to the real-time tracking of multiple genes across varying temporal scales. Finally, by monitoring the dynamics of *CDKN1A*, *PCNA*, and *VCL* across distinct transcriptional states, we reveal that the effects of transcriptional modulation on chromatin motion is gene-dependent and does not necessarily induce significant alterations in local chromatin mobility. This approach opens new avenues for dissecting the interplay between genome organization and gene regulation and offers a generalizable tool for investigating chromatin dynamics in development, disease, and response to environmental cues.

## Results

### Multivalent dSaCas9-ArrayG/N label enables long-term, selective, and photostable chromatin tracking

To track chromatin loci with high spatial and temporal resolution over extended durations, we developed a replenishable labeling system based on a fusion of dSaCas9 and a multivalent ArrayG/N module. This construct combines 8x or 16x GFP-binding nanobodies with 2 or 4 importin β–binding (IBB) domains to support both robust signal amplification and nuclear import (Fig. 1a). We selected dSaCas9 over the more commonly used dSpCas9 due to its smaller size (∼3.2 kb vs. ∼4.2 kb), which improves delivery efficiency and allows greater flexibility for modular design. In addition, dSaCas9 requires a longer PAM sequence (NNGRRT), which reduces off-target binding and enhances targeting specificity^53^. When expressed alongside monomeric wild-type GFP-GB1 in U2OS cells, dSaCas9-ArrayG/N formed distinct, trackable puncta, indicating efficient chromatin binding (Fig. 1b, Supplementary Fig. 1). To confirm target specificity, we tracked the dynamics of labeled loci in the presence of no sgRNA, a non-targeting sgRNA, or a sgRNA targeting repetitive Alu elements. Only the targeting sgRNA induced prolonged chromatin binding (Fig. 1c,d), demonstrating the high specificity of the system.

**Figure 1:**
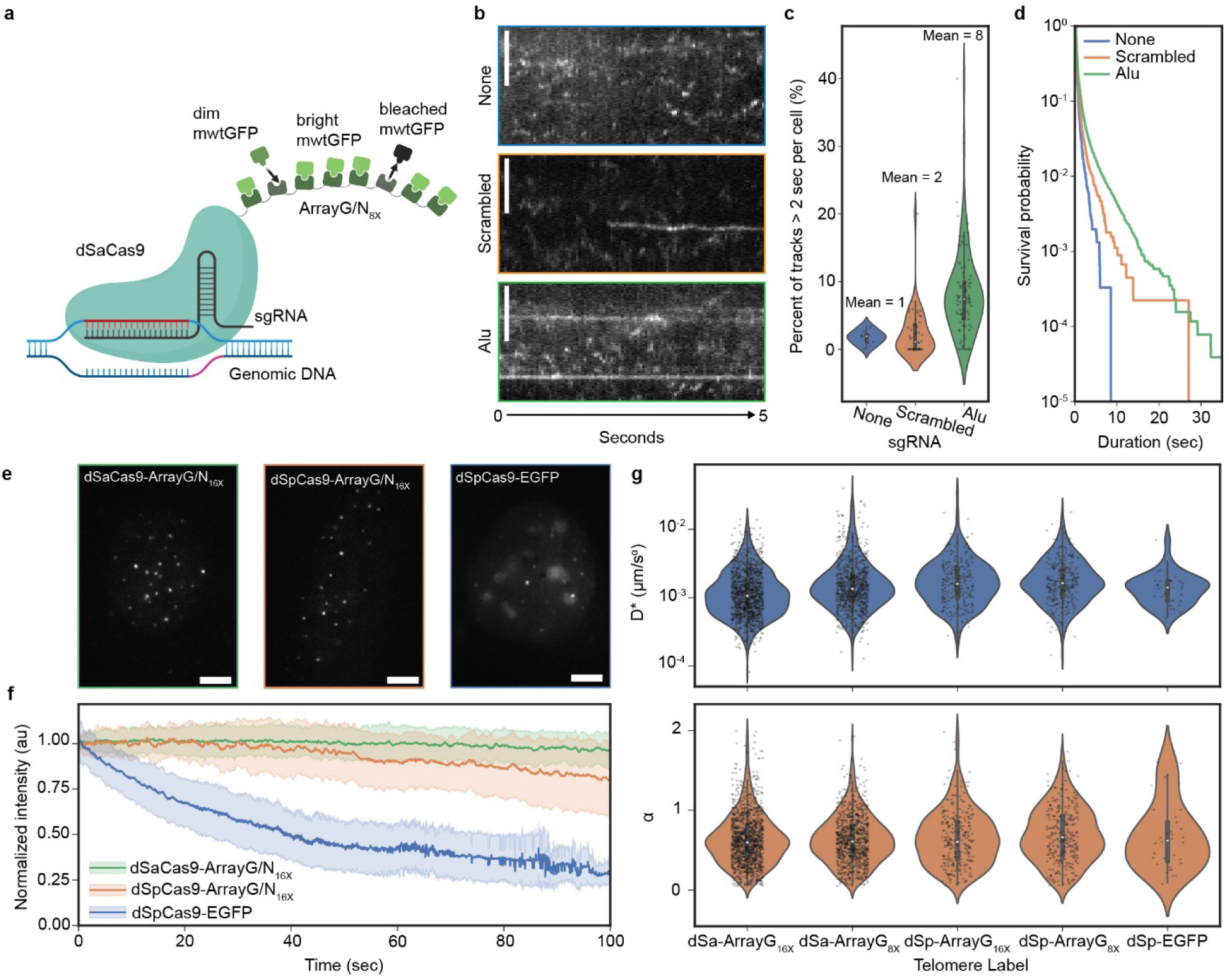
Multivalent dSaCas9-ArrayG/N label enables long-term, selective, and photostable chromatin tracking. **a** Simplified cartoon of dSaCas9-ArrayG/N_8X_ (not shown to scale). **b** Kymographs of U2OS cells expressing dSaCas9-ArrayG/N_16X_ (*top*) without sgRNA (None), (*middle*) with a non-targeting/scrambled sgRNA (Scrambled), and (*bottom*) with Alu-targeting sgRNA (Alu). Intensity normalized independently for each image. Scale bars are 1 µm. **c** Violin plots of the proportion of tracks per cell greater than 2 seconds for each sgRNA condition (*n* = 12, 25, and 87 cells for None, Scrambled, and Alu, respectively). **d** Survival probability of trajectories for each sgRNA condition (*n* = 29,500, 52,614, and 400,063 trajectories in 12, 25, and 87 cells for None, Scrambled, and Alu, respectively). **e** Representative images (single focal planes) of telomeres labeled with dSaCas9-ArrayG/N_16X_, dSpCas9-ArrayG/N_16X_, and dSpCas9-EGFP. Intensity normalized independently for each image. Scale bars are 5 µm. **f** Intensity decay of telomeres labeled by the indicated dCas9 tags and imaged at 20 Hz (data shows mean ± std, *n* = 228, 78, and 103 trajectories in 10 cells each for dSaCas9-ArrayG/N_16X_, dSpCas9-ArrayG/N_16X_, and dSpCas9-EGFP, respectively). **g** Mean squared displacement analysis of telomeres labeled with the indicated dCas9 tags and tracked at 20 Hz, shown as violin plots of the fitted (*top*) effective diffusion coefficients, D*, and (*bottom*) anomalous exponents, α (*n* = 1276, 884, 265, 294, 34 trajectories across 78, 57, 40, 40, 30 cells, from left to right).

We next benchmarked dSaCas9-ArrayG/N against both dSpCas9-ArrayG/N and conventional dSpCas9-EGFP by targeting telomeres and imaging under continuous illumination. While dSpCas9-EGFP signals rapidly decayed due to photobleaching, dSaCas9-ArrayG/N and dSpCas9-ArrayG/N maintained stable fluorescence, consistent with dynamic fluorophore exchange (Fig. 1e,f).

To assess whether increasing the array size alters chromatin dynamics, we compared the trajectories of telomeres labeled with 8x or 16x arrays fused to dSaCas9 or dSpCas9, as well as with the conventional dSpCas9-EGFP. Mean squared displacement (MSD) analysis revealed similar telomere dynamics across all dCas9 labels used. There were slight differences in D*, which were not correlated with label size or Cas9 ortholog (Fig. 1g, Supplementary Fig. 2, Supplementary Table 1). Thus, the dCas9-ArrayG/N system provides a versatile and minimally invasive platform for long-term, high-fidelity chromatin tracking.

### dSaCas9-ArrayG/N enables efficient labeling and quantitative tracking of repetitive chromatin loci

Having demonstrated that dSaCas9-ArrayG/N provides high specificity and sustained signal, we next tested its capacity to label endogenously repetitive regions of the genome. These loci offer a unique opportunity to evaluate signal amplification, labeling uniformity, and the system’s suitability for comparing chromatin dynamics. To this end, we identified 17 previously uncharacterized loci containing clusters of at least 10 dSaCas9 target sites, spaced 55–1530 bp apart within 3.5–50.2 kb windows, and included 3 additional loci previously reported in the literature (Fig. 2a, Supplementary Table 2). All 20 regions exhibited bright nuclear puncta upon targeting, consistent with robust and specific labeling by dSaCas9-ArrayG/N (Fig. 2b, Supplementary Fig. 3).

**Figure 2:**
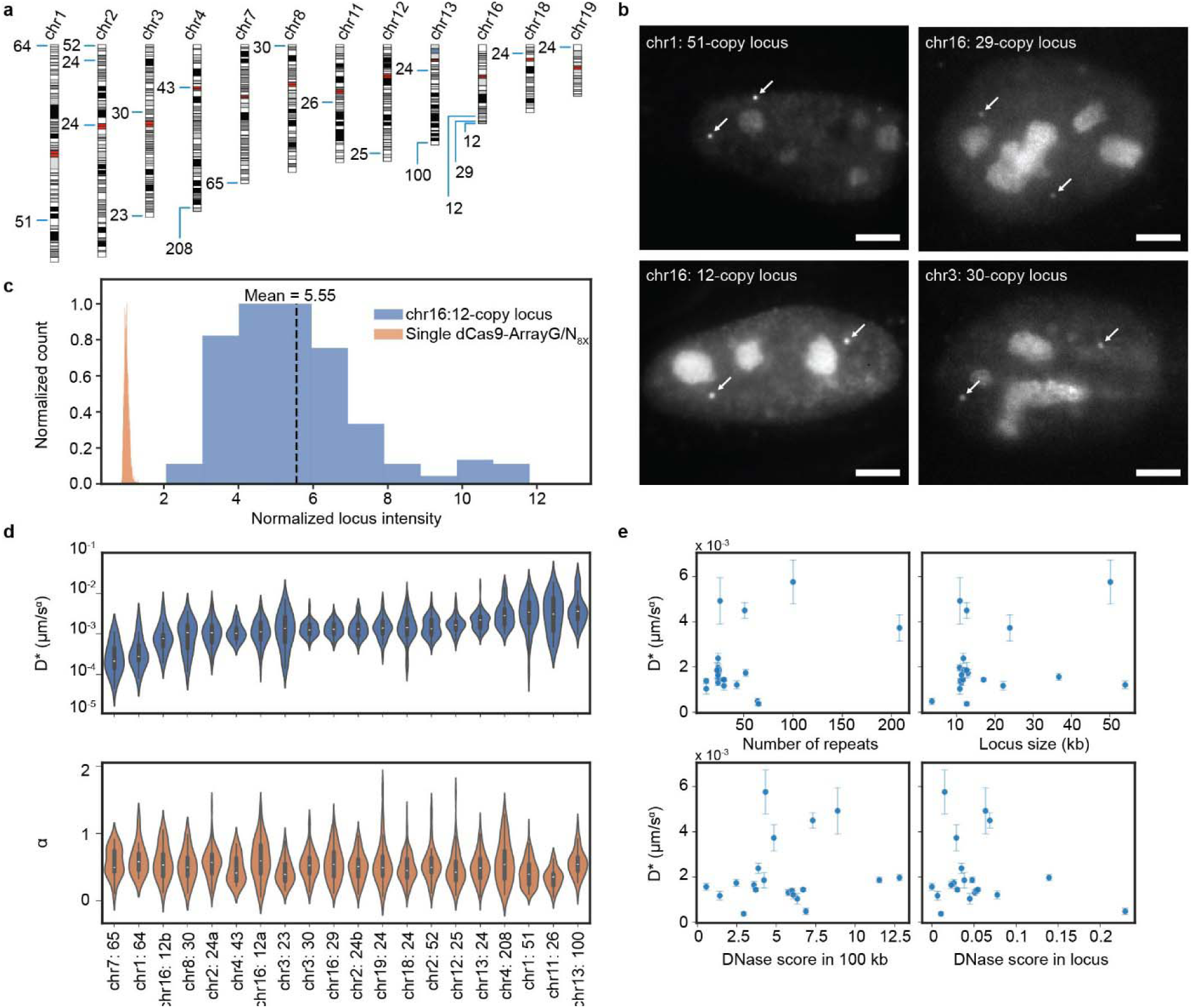
dSaCas9-ArrayG/N enables efficient labeling and quantitative tracking of repetitive chromatin loci. **a** Ideogram of selected repetitive loci harboring SaCas9 targets. The chromosome number is indicated at the top, and the annotations show the number of repeats (target sites) at that locus. **b** Representative images (single-plane snapshots) of the indicated repetitive locus labeled with dSaCas9-ArrayG/N_8X_. Arrows point to loci. Intensity normalized independently for each image. Scale bars are 5 µm. **c** Intensity histogram of single dSaCas9-ArrayG/N_8X_ and a 12-copy locus on chromosome 16 labeled by dSaCas9-ArrayG/N_8X_, in units of the mean intensity of single dSaCas9-ArrayG/N_8X_ (*n* = 48 spots across 24 cells for chr16: 12-copy locus and 473 spots across 6 cells for single dSaCas9-ArrayG/N_8X_). **d** Mean squared displacement analysis of all 20 repetitive loci from **a** tracked at 5 Hz, shown as violin plots of the fitted (*top*) effective diffusion coefficients, D*, and (*bottom*) anomalous exponents, α (*n* > 13 trajectories for each locus; details in Supplementary Table 2). The suffix “a” and “b” are used when there are multiple loci with the same copy-number on the same chromosome, where “a” indicates the first in genomic coordinate space. All loci were labeled by dSaCas9-ArrayG/N_8X_ except chr1: 64, chr7: 65, and chr3: 23, which were labeled by dSaCas9-ArrayG/N_16X_. **e** Plots of the effective diffusion coefficient for each locus plotted against the indicated characteristics of that locus (data shows mean ± SEM for *n* > 13 trajectories for each locus). No correlation (r^2^ < 0.17) was found for any plot.

To quantify labeling efficiency, we compared the fluorescence intensity of a 12-repeat locus on chromosome 16 to the average intensity of single dSaCas9-ArrayG/N puncta. This analysis indicated ∼50% occupancy, suggesting that as few as six bound complexes are sufficient to achieve high SBR at repetitive loci (Fig. 2c).

Tracking across all labeled loci revealed variability in effective diffusion coefficients D*, while the anomalous exponent α remained relatively constant (Fig. 2d, Supplementary Table 2). Surprisingly, D* did not correlate with locus size, repeat count, or chromatin accessibility (Fig. 2e), suggesting that additional contextual or epigenetic factors may govern local chromatin dynamics. Although replicate variability was observed (Supplementary Fig. 4), it was substantially lower than the variation across different loci, underscoring the biological heterogeneity in chromatin motion.

### Multiplexed polycistronic sgRNA expression enables robust labeling of single-copy loci using dSaCas9-ArrayG/N

While repetitive genomic loci offer convenient targets for chromatin tracking due to inherent fluorescence signal amplification, most biologically relevant loci are non-repetitive and require numerous sgRNAs for effective visualization. However, existing strategies such as tandem U6-driven expression suffer from poor viral delivery due to their large size and impose sequence constraints; each sgRNA must begin with a 5′ “G”, which limits guide design flexibility (Fig. 3a).

**Figure 3:**
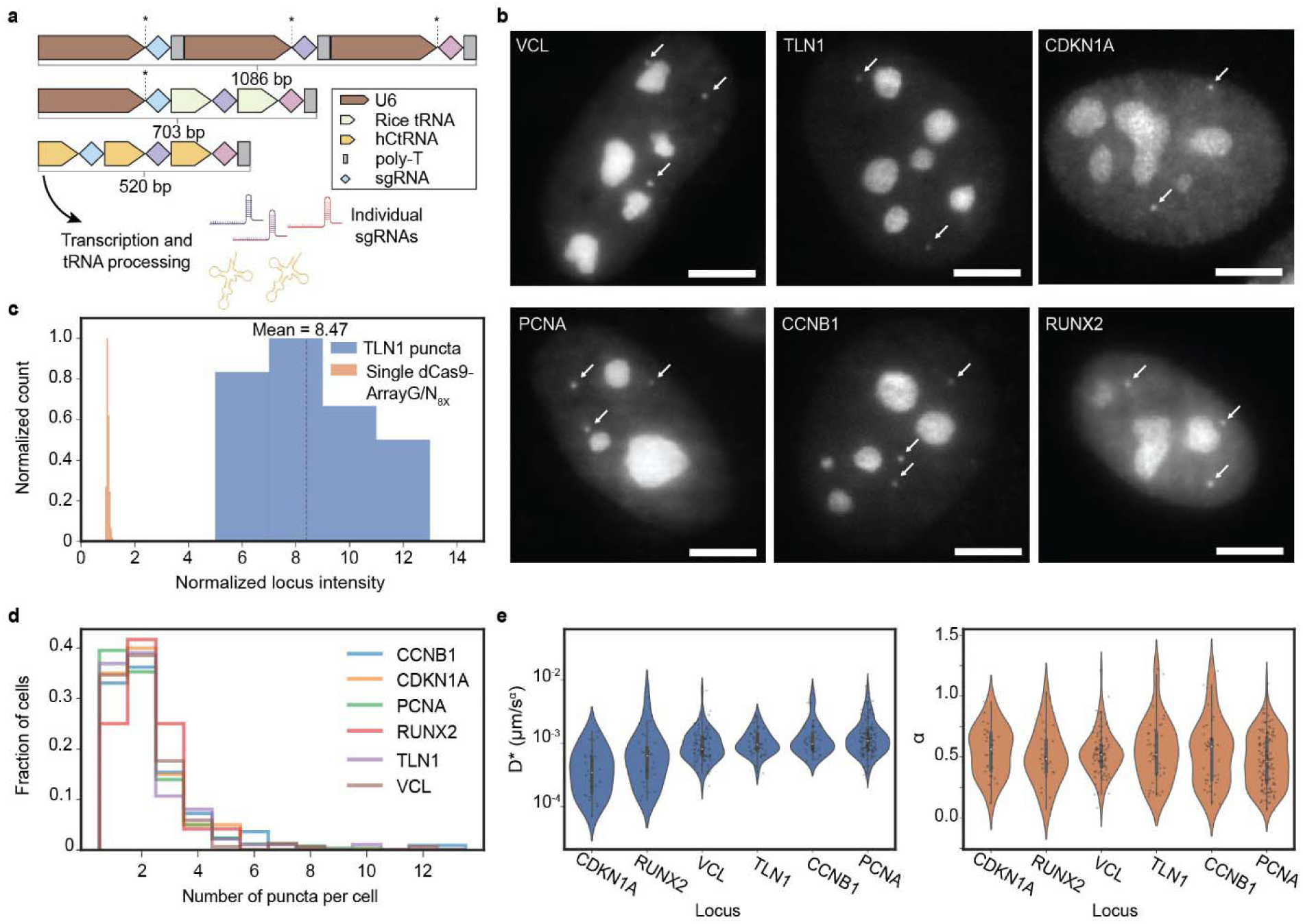
Multiplexed polycistronic sgRNA expression enables robust labeling of single-copy loci using dSaCas9-ArrayG/N. **a** Schematic of the three ways used to express multiple unique sgRNAs from a single plasmid. “*” is used to indicate where a sgRNA must begin with the “G” nucleotide for efficient expression by the U6 promoter. **b** Representative images (single-plane snapshots) of the indicated gene labeled with dSaCas9-ArrayG/N_8X_ and 10 sgRNAs driven from PTG construct (*VCL, TLN1, PCNA, CCNB1*) or dSaCas9-ArrayG/N_16X_ and 8 sgRNAs driven from DAP construct (*CDKN1A, RUNX2*). Arrows point to loci. Intensity normalized independently for each image. Scale bars are 5 µm. **c** Intensity histogram of single dSaCas9-ArrayG/N_8X_ and *TLN1* labeled by dSaCas9-ArrayG/N_8X_ in units of the mean intensity of single dSaCas9-ArrayG/N_8X_ (*n* = 18 spots across 12 cells for *TLN1* and 656 spots across 26 cells for single dSaCas9-ArrayG/N_8X_). **d** Spot count histograms for each labeled gene in cells with at least 1 spot. Spots were counted in max-intensity projections of z-stacks covering the full nucleus (*n* = 221, 20, 258, 20, 153, and 187 cells for *CCNB1*, *CDKN1A*, *PCNA*, *RUNX2*, *TLN1*, and *VCL*, respectively). **e** Diffusion analysis of all six genes tracked at 5 Hz, shown as violin plots of the fitted (*left*) effective diffusion coefficients, D*, and (*right*) anomalous exponents, α (*n* = 39, 26, 111, 23, 41, 81 trajectories across 30, 20, 78, 21, 34, 43 cells for *CCNB1*, *CDKN1A*, *PCNA*, *RUNX2*, *TLN1*, and *VCL*, respectively).

To address these challenges, we first tested a conventional construct encoding six individually driven U6 sgRNAs targeting the *VCL* locus. While this approach successfully generated nuclear puncta (Supplementary Fig. 5), its size and rigidity hindered scalability and packaging efficiency.

To overcome these limitations, we implemented two compact, polycistronic sgRNA strategies: the tRNA–sgRNA (PTG) and drive-and-process (DAP) systems^54,55^. In the PTG approach, rice tRNA sequences are interleaved between sgRNAs, enabling expression of multiple guides from a single U6 promoter and lifting the 5′ “G” constraint. In the DAP system, a human cysteine tRNA (hCtRNA) acts as both the promoter and processing element, allowing sgRNA arrays to be transcribed without a U6 promoter. We utilized PTG– and DAP-based polycistronic constructs to express multiple sgRNAs from a single transcript. This strategy simplifies construct design, enhances sgRNA delivery efficiency, and avoids the need for *in vitro* transcription or chemical modification, offering a cost-effective and scalable solution for labeling diverse genomic regions. Importantly, these systems provide a fully genetically encoded means of delivering multiplexed sgRNAs, eliminating the need for synthetic RNA delivery or pooled transfections.

To validate these systems, we expressed a telomere-targeting array using the DAP system and observed robust nuclear puncta even from the eighth sgRNA position, indicating efficient transcript processing and functional sgRNA output (Supplementary Fig. 6). We then applied PTG and DAP multiplexing to target six non-repetitive endogenous genes (*PCNA*, *CCNB1*, *TLN1*, *VCL*, *RUNX2*, and *CDKN1A*), designing 8–10 sgRNAs per locus that flanked ∼2–4 kb regions upstream or downstream of each gene (Supplementary Fig. 7, Supplementary Table 3).

Expression of these constructs in cells stably expressing dSaCas9-ArrayG/N yielded bright nuclear puncta across all loci (Fig. 3b). Quantification of *TLN1* puncta intensities revealed signal levels consistent with the number of sgRNAs used, supporting the use of multi-guide arrays to amplify chromatin labeling (Fig. 3c). Max-intensity z-projections showed 1–3 puncta per cell, consistent with the trisomic karyotype of U2OS cells (Fig. 3d). Tracking of these loci revealed chromatin dynamics within the range observed at repetitive elements, with modest heterogeneity between genes (Fig. 3e, Supplementary Fig. 8, Supplementary Table 4).

The successful labeling of six distinct endogenous loci across diverse genomic contexts demonstrates the generalizability of this approach and supports its use as a scalable framework for single-locus chromatin tracking. Furthermore, as a fully genetically encoded system, our approach is compatible with stable cell line generation and viral delivery, making it well-suited for longitudinal studies of chromatin architecture during differentiation, reprogramming, or disease progression.

### Chromatin motion at endogenous gene loci is not necessarily dependent on transcriptional activity

Whether transcription directly influences local chromatin mobility remains an open question. Prior studies have reported correlations between gene activity and chromatin dynamics, but many relied on synthetic reporters or indirect ensemble measurements that lack locus-specific resolution^20,21,35–37,56^. To address this, we utilized MS2-MCP RNA labeling^57,58^, enabling a dual-color genetically encoded system to directly measure the relationship between transcriptional activity and chromatin motion at endogenous, non-repetitive loci.

We first asked whether steady-state transcription levels correlate with chromatin mobility across different genes. Plotting the average diffusion coefficient (D*) of six labeled endogenous loci against their reported expression levels in U2OS cells (sourced from the Human Protein Atlas^59^) revealed no direct correlation (Supplementary Fig. 9). This initial finding suggests that basal gene expression does not predict chromatin dynamics at these loci.

To test the transcription-dynamics relationship more directly, we perturbed transcription pharmacologically and tracked the resulting chromatin motion. Inhibition of transcription initiation using triptolide did not significantly alter the high-frequency (20 Hz) dynamics of the *CDKN1A* or *VCL* loci, but it did increase the average diffusion coefficient of *PCNA* loci (Fig. 4a, Supplementary Table 5). We then activated *CDKN1A* expression with Nutlin-3a, a small molecule that stabilizes p53 and increases the fraction of transcriptionally active cells from ∼20% to ∼50% within tens of minutes (Supplementary Fig. 10)^60^. Despite robust transcriptional activation, locus tracking revealed no significant change in mobility between untreated and Nutlin-3a-treated cells over a 15–60 min window (Fig. 4b, Supplementary Table 5).

**Figure 4:**
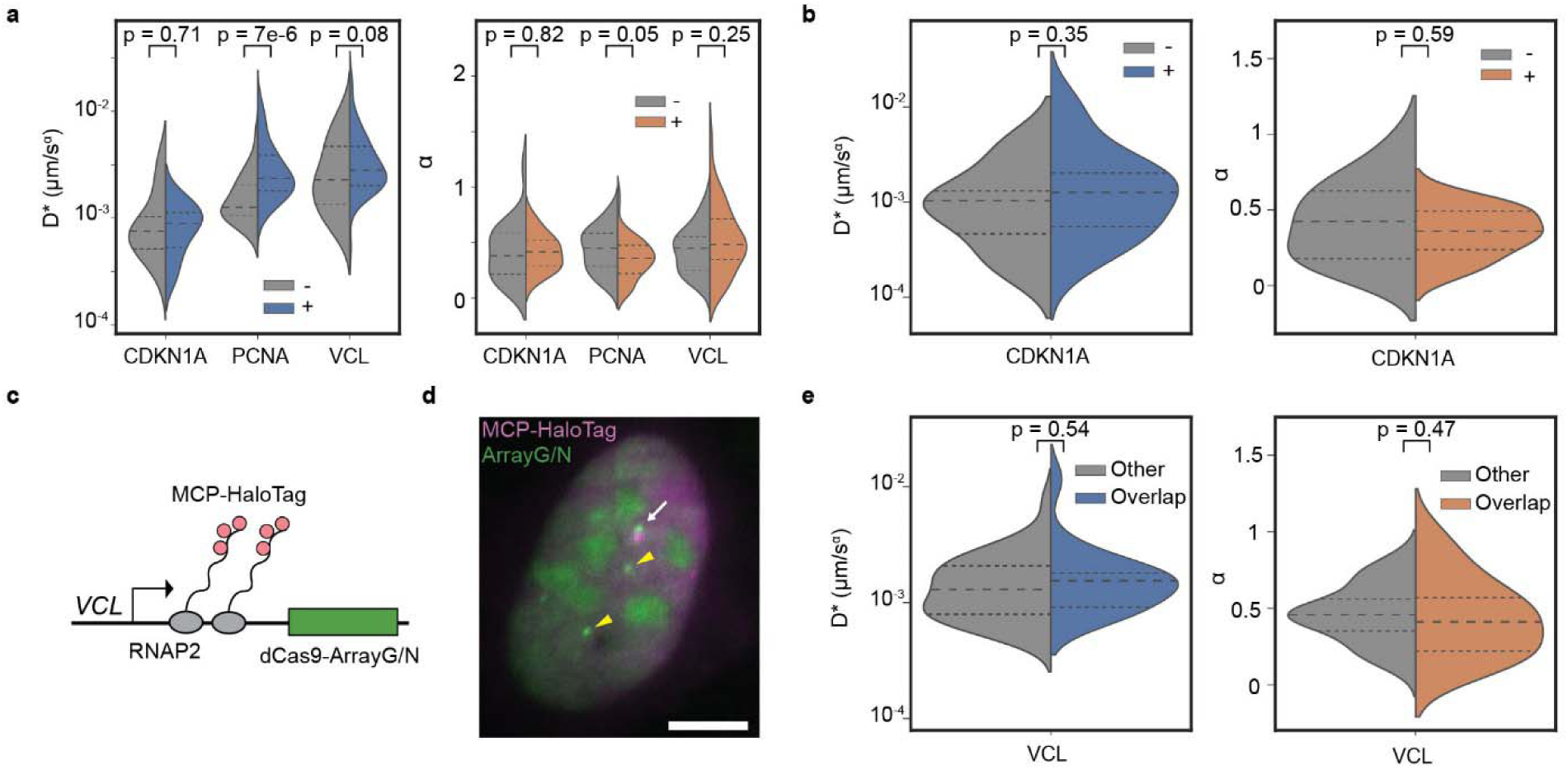
Chromatin motion at endogenous gene loci is not necessarily dependent on transcriptional activity. **a** Mean squared displacement analysis of *CDKN1A*, *PCNA*, and *VCL* without (−) and with (+) triptolide treatment, tracked at 20 Hz, shown as split violin plots of the fitted (*left*) effective diffusion coefficients, D*, and (*right*) anomalous exponents, α (*CDKN1A: n* = 22 and 18 trajectories across 19 and 13 cells for untreated and treated, respectively; *PCNA*: *n* = 31 and 69 trajectories across 29 and 56 cells for untreated and treated, respectively; *VCL: n* = 35 and 50 trajectories across 33 and 47 cells for untreated and treated, respectively). **b** Mean squared displacement analysis of *CDKN1A* without (−) and with (+) Nutlin-3a treatment, tracked at 20 Hz, shown as split violin plots of the fitted (*left*) effective diffusion coefficients and (*right*) anomalous exponents (*n* = 15 and 23 trajectories across 15 and 21 cells for untreated and treated, respectively). **c** Schematic of two-color labeling: the nascent RNA is labeled via an MS2 insertion into *VCL* 3’ UTR and expression of MCP-HaloTag; the chromatin downstream of the 3’ UTR is labeled by dSaCas9-ArrayG/N_16X_. **d** Representative two-color image of the nascent transcription spot (NTS) overlapping with the dSaCas9-ArrayG/N-labeled *VCL*. The white arrow indicates overlapping NTS and chromatin spot whereas the yellow triangles indicate chromatin spots without a corresponding NTS (“Other”). Scale bar is 5 µm. **e** Mean squared displacement analysis of *VCL* loci overlapping with the NTS (i.e., actively transcribing) and all other loci, tracked at 20 Hz, shown as split violin plots of the fitted (*left*) effective diffusion coefficients and (*right*) anomalous exponents (*n* = 40 and 20 trajectories across 30 and 20 cells for other and overlap, respectively). All p-values are the results of two-sided Mann-Whitney U tests.

To resolve potential heterogeneity at the single-cell level, we integrated 24 MS2 stem loops into the 3′ UTR of *VCL* and labeled nascent transcripts using MCP-HaloTag conjugated to a JF646 dye^61^. This allowed direct visualization of transcriptional activity (nascent transcription spots, NTS) alongside the chromatin locus via dSaCas9-ArrayG/N (Fig. 4c,d). We then classified loci as transcriptionally active based on spatial overlap with the NTS and measured their dynamics. Strikingly, transcriptionally active *VCL* loci exhibited no significant difference in diffusion behavior compared to other loci, which could be active or inactive, within the same nucleus (Fig. 4e, Supplementary Table 5). These results suggest that transcriptional activity alone does not significantly modulate the local physical dynamics of chromatin at the *VCL* locus, at least on the spatial and temporal scales probed here.

Our findings are consistent with recent high-precision tracking studies by Mazzocca *et al*., who used MINFLUX imaging to monitor chromatin motion over seven orders of magnitude in time and found that chromatin is highly subdiffusive and only weakly affected by transcriptional perturbation^62^. While Mazzocca *et al.* focused on global chromatin and engineered reporters, our data extend this conclusion to multiple endogenous gene loci using a genetically encoded, dual-color live-cell system. Our results also complement recent experiments by Chu *et al.*, which demonstrated that transcription-dependent changes in chromatin motion are gene dependent and governed by local chromatin compaction^35^. Together, these complementary approaches suggest a growing consensus: chromatin mobility is governed more by nuclear architecture and polymer constraints than by real-time transcriptional state.

## Discussion

We present a genetically encoded, high-resolution platform for labeling and tracking non-repetitive genomic loci in living cells that overcomes several longstanding limitations of existing approaches. This system combines a photostable dSaCas9-ArrayG/N fusion with compact polycistronic sgRNA expression, enabling (1) full genetic encodability and compatibility with long-term or *in vivo* studies, (2) high-precision tracking sustained over thousands of frames, (3) efficient and scalable delivery of multiple sgRNAs per locus, (4) broad targeting flexibility for both repetitive and single-copy genomic regions, (5) orthogonal compatibility with other tagging systems such as dSpCas9 and FROS for multimodal imaging, and (6) straightforward implementation using standard molecular and imaging tools. Using this platform, we labeled and tracked 20 repetitive regions and six endogenous, non-repetitive gene-adjacent loci in live cells. Chromatin motion exhibited modest gene-specific variability and pronounced differences between repetitive regions. Notably, these patterns did not correlate with chromatin accessibility in a simple linear fashion, indicating that additional factors, such as local chromatin topology or epigenetic modifications, likely play a role in shaping locus-specific dynamics. Finally, we found that global transcription inhibition had a gene-specific effect on chromatin dynamics, and by combining our approach with chemically induced transcriptional activation and with the MS2-MCP reporter system, we observed that chromatin mobility at *CDKN1A* and *VCL* loci remained largely unaffected by transcriptional state, supporting a growing view that transcription and local chromatin motion could be mechanistically decoupled and that the relationship is context dependent.

In this study, we labeled genomic regions located several kilobases upstream or downstream of genes to avoid potential interference by dCas9 binding with endogenous regulatory activity. While this conservative strategy is designed to preserve native chromatin behavior, the low number of dCas9 complexes required for effective labeling in our system increases the feasibility of direct targeting of gene bodies, promoters, or distal regulatory elements, such as enhancers. Achieving ∼80% labeling efficiency at the *TLN1* locus with just 10 sgRNAs underscores the potential for sparse guide designs. This capability enables real-time interrogation of regulatory elements involved in transcriptional control, cell fate transitions, and disease-linked chromatin dysregulation. Defining how the proximity of labeling affects chromatin mobility and function will be critical for enabling high-resolution tracking without perturbation. Applications such as allele-specific labeling or multiplexed regulatory element tracking will benefit from rigorous optimization and validation pipelines tailored to the chromatin context of each locus.

Our findings contribute to a growing body of evidence that the relationship between transcription and chromatin dynamics is highly gene– and context-dependent. While some studies have reported increased mobility upon transcriptional activation^35,37^, others have found decreased motion^20,21,36,56^ or no measurable change^20,21^. A recent study revealed that only loci situated in less compact chromatin exhibited increased dynamics upon activation^35^, highlighting the influence of the local chromatin environment. Additionally, three-color imaging of the promoter of *Sox2*, its enhancer, and transcription revealed homogenization of enhancer and promoter dynamics upon transcriptional activation – the enhancer became more constrained, whereas the promoter demonstrated an increased dynamics^63^. These diverse outcomes underscore the need for high-resolution, locus-specific, and temporally resolved measurements in native genomic contexts. The flexibility of our labeling platform, enabling precise tracking of endogenous loci under defined transcriptional or epigenetic states, makes it well-suited to address these open questions. By allowing simultaneous imaging of chromatin motion and transcriptional activity at specific sites, our approach can help disentangle the interplay between gene regulation, nuclear architecture, and chromatin mechanics. Our approach can also easily be extended for 3D tracking and imaging using point spread function engineering and with contrast improvement through light sheet illumination^52,64–72^.

The flexibility and genetic encodability of our approach make it readily adaptable for tracking the dynamics of additional genes, regulatory elements, or structural features across diverse cell types and conditions. Its complete genetic encodability makes it especially useful in settings where *in vitro* delivery of preassembled dCas9–sgRNA complexes is impractical, such as in developmental models, long-term imaging experiments, or lineage-tracing studies. The ability to express all components from genome-integrated DNA cassettes allows stable, controllable, and minimally invasive labeling in living systems. Beyond single-locus tracking, our method is compatible with orthogonal tagging systems like dSpCas9 or FROS, enabling two-color imaging of chromatin interactions, for example, between enhancers and promoters. It also supports multimodal experiments, such as simultaneously tracking a transcription factor (TF) and its target gene, to directly observe TF-chromatin interactions in live cells. Together, these capabilities make our platform a powerful tool for dissecting genome organization and its regulation in space and time.

The versatility and modularity of this system render it well-suited for a wide array of future applications. Its compact, genetically encoded design enables high-throughput or multiplexed tracking of genomic loci across different conditions, making it compatible with large-scale studies of genome regulation. The ability to express all components stably also enables integration with perturbation platforms such as CRISPR screens or Perturb-seq^73–75^, allowing functional dissection of chromatin behavior alongside gene expression. In developmental and disease models, where repeated *in vitro* delivery is impractical, this system offers a minimally invasive, long-term labeling strategy compatible with studying chromatin dynamics in stem cell–derived or patient-derived systems^76,77^. Additionally, the high-resolution trajectory data produced by our method provides a rich framework for testing physical models of chromatin organization, including constrained polymer dynamics and nuclear compartmentalization^78^. Finally, the system’s photostability and temporal control make it ideally suited for longitudinal studies of chromatin remodeling, such as during differentiation, senescence, or stress response^3,6^. Together, these capabilities position our platform as a versatile tool for probing genome function and dynamics across scales, contexts, and experimental modalities.

## Methods

### Cell culture

Human osteosarcoma (U2OS) cells (ATCC, U-2 OS HTB-96) and U2OS p21-MS2 MCP-EGFP cells (gift from Dr. Robert Singer^79^) were cultured in Dulbecco’s Modified Eagle’s Medium (DMEM) without phenol red (Gibco, 21063029) supplemented with 10% (v/v) fetal bovine serum (FBS) (Gibco, A5669701) and 1 mM sodium pyruvate (Gibco, 11360070) at 37°C and 5% CO_2_. HEK293T cells (ATCC, CRL-3216) were cultured in DMEM (ATCC 30-2002) supplemented with 10% (v/v) FBS (Gibco, A5669701) and 2 mM L-glutamine (ATCC 30-2214). Cells stably expressing dSaCas9-ArrayG/N and mwtGFP were cultured in media also containing 250 μg/ml hygromycin (InvivoGen, ant-hg-1) and 10 μg/ml blasticidin (InvivoGen, ant-bl-05). Cells additionally stably expressing sgRNA were cultured in media also supplemented with 1 μg/ml puromycin (InvivoGen, ant-pr-1). 48 hours before imaging, media was replaced with antibiotic-supplemented complete media containing 1 μg/ml doxycycline (Sigma, D9891) and 1x cumate (System Biosciences, QM100A-1) to induce expression of dCas9-ArrayG/N and mwtGFP, respectively.

### Cell line generation

To generate U2OS cells stably expressing dCas9-ArrayG/N and mwtGFP, U2OS cells were transfected with plasmids pRPG101, pRPG102, and piggyBac transposase (see Supplementary Table 6) using Lipofectamine 3000 (Thermo Fisher, L3000015) following the manufacturer’s protocols in a 6-well plate. A plasmid ratio of 2:2:1 was used (see Supplementary Table 7). Two days after transfection, media was replaced with media supplemented with 250 μg/ml hygromycin and 10 μg/ml blasticidin and changed every other day until selection was complete. 3^rd^ generation lentivirus was produced via transfection of HEK293T cells with plasmids pMDLG/pRRe (Addgene #12251, gift from Dr. Didier Trono), pMD2.G (Addgene #12259, gift from Dr. Didier Trono), and pRSV-Rev (Addgene #12253, gift from Dr. Didier Trono) (Supplementary Table 6) and the corresponding transfer plasmid using Lipofectamine 3000 (see Supplementary Table 7 for plasmid ratios). Supernatant was harvested 24 and 48 hours post transfection, filtered through a 0.45 μm filter, and concentrated using PEG-it viral precipitation kit (System Biosciences, LV810A-1) following the manufacturer’s protocols. To generate cells stably expressing the desired sgRNA, U2OS cells stably expressing dSaCas9-ArrayG/N were reverse transduced with varying levels of concentrated virus in a 6-well plate and selected with 1 μg/ml puromycin starting two days later. To generate the U2OS-VCL-MS2 cell line, cells were transfected with one plasmid encoding SpCas9 and sgRNA (pX330, Addgene #42230, a gift from Feng Zhang) targeting just upstream of the *VCL* stop codon, and a synthesized homology donor consisting of LHA-linker-mCherry-P2A-BlastR-24xMS2-RHA. Successfully edited cells were selected with blasticidin and confirmed by imaging endogenous VCL-mCherry in its correct location and visualizing nascent transcription sites after transduction with MCP-HaloTag-GB1 (pRPG200 in Supplementary Table 6). Then, dSaCas9-ArrayG/N and mwGFP-GB1 were introduced as described above for the U2OS cell line to generate the dual-color cell line.

### Plasmids and cloning

All plasmids used in this study are listed in Supplementary Table 6. We used typical cloning protocols for all constructs, using either Gibson assembly (NEBuilder® HiFi DNA Assembly Master Mix, E2621S) or Golden Gate assembly. Miniprepped plasmids (QIAGEN, 12123) were checked via restriction digest and full-plasmid sequencing (Plasmidsaurus). Plasmids for transfections were maxiprepped (QIAGEN, 12162) and sequenced again. All primers were ordered from Integrated DNA Technologies, Inc. (IDT) as ssDNA oligonucleotides.

For DAP array cloning: the vector and fragment template plasmids were obtained (hCtRNA_VT (Addgene #186716) and hCtRNA_FT (Addgene #186715), gifts from Dr. Xue Gao) and the sgRNA scaffold was replaced with the optimized scaffold for dSaCas9^80^ by cloning the dSaCas9-optimized scaffold, which was ordered as dsDNA in a gBlock (IDT), into the PCR-amplified template (either hCtRNA_VT or hCtRNA_FT) using NEBuilder HiFi Assembly (NEB, E2621S). An analogous approach was used to replace the sgRNA scaffold with one optimized for dSpCas9^11^ imaging to be used in the telomere labeling experiment with DAP-driven sgRNA (Supplementary Fig. 6). To generate a DAP array, the protocol from the original paper^55^ was followed with some modifications. Briefly, primers were designed to amplify 1 vector template and *N*-1 fragment templates, for *N* sgRNAs, containing about half of the sgRNA spacer sequence and BsaI cut sites, such that the overhangs were compatible with Golden Gate assembly with NEBuilder BsaI-HFv2 (NEB, E1601S). The PCR details are shown in Supplementary Table 8. PCR amplicons (1 vector template, *N-*1 fragment templates) of the correct size were excised from a gel, purified, used for Golden Gate assembly with NEBuilder BsaI-HFv2 (NEB, E1601S) following the manufacturer’s protocols, and 2 μl of the product was transformed into NEB Stable Competent *E. coli* (High Efficiency) cells (NEB, C3040H) following the manufacturer’s protocol (see Supplementary Fig. 7 for example gels and schematic of the cloning process). Miniprepped plasmids were checked for size by restriction digest with BamHI-HF (NEB, R3136S) and XhoI (NEB, R0146S) and then confirmed by full-plasmid sequencing (Plasmidsaurus). To generate lentiviral transfer vectors, the DAP array was digested with BamHI-HF and XhoI, the correct band excised from a gel, and inserted into a PCR-amplified 3^rd^ generation lentiviral transfer vector containing a puromycin resistance cassette (pAR220 in Supplementary Table 6, which was derived from pLH-spsgRNA2 (Addgene #64114, gift from Dr. Thoru Pederson)), containing overhangs matching the ends of the DAP array using NEBuilder HiFi Assembly (NEB, E2621S).

For PTG array cloning: tRNA-sgRNA constructs were assembled using hierarchical Golden Gate assembly. To perform Golden Gate assembly, a lentiviral transfer vector for SaCas9 sgRNA expression was modified to harbor BsaI cut sites after a U6 promoter driving a SaCas9 scaffold-tRNA cassette: U6-BsaI-BsaI-scaffold-tRNA (pRPG100 in Supplementary Table 6). The tRNA-sgRNA 10x array was first assembled as two pieces using Golden Gate assembly, each piece containing tRNA-sgRNA 5x arrays. Then the two pieces were assembled using restriction enzyme sites digestion-ligation. The first step was to synthesize the individual sgRNA-tRNA pieces. Each sgRNA consists of two pieces: a unique 20 bp spacer and the 76 bp optimized SaCas9 scaffold. Using scaffold-tRNA (pRPG100) as the template, PCR was performed with primers including the unique spacer and BsaI cut sites such that the fragment 1-5 and 6-10 could be assembled in a Golden Gate assembly reaction into the lentiviral transfer vector. One extra bp (guanine) was inserted before the start of the first fragment to enable the functionality of the U6 promoter, and polyT (tttttt) was added to the last fragment to terminate the transcription driven by the U6 promoter. The tRNA module from the last fragment was removed to avoid extra base pairs. To ligate the two 5x arrays together, an extra SalI site was inserted at the end of the 5^th^ fragment and beginning of the 6^th^ fragment. After the Golden Gate assembly, the two resulting arrays are: (1) 5’…-U6-5xArray-SalI-BamHI…3’, and (2) 5’…-SalI-5xArray-BamHI…3’. Finally, the plasmids with the two 5x arrays were digested with BamHI-HF and SalI (NEB, R0138S), gel purified and ligated to create the 10x array.

### sgRNA design

5 kb regions adjacent to a gene of interest were chosen and input to the online sgRNA design tool CHOPCHOP^81^ to generate candidate sgRNAs for imaging. Then, sgRNAs were manually chosen such that 6-10 sgRNAs were spaced ∼ 500 bp apart, corresponding to a targeted region of 3-5 kb. This spacing was chosen to minimize spatial interference between neighboring dCas9-ArrayG/Ns and to not disrupt the chromatin at a concentrated region. An example pipeline is demonstrated in Supplementary Fig. 7.

### Cell imaging

For transient sgRNA expression via lipofectamine (for Supplementary Figs. 5 and 6), the protocol was as follows: three days before imaging, 300 μl [5 x 10^4^ cells/ml] of cells were plated in 8-well glass bottom chambers (ibidi, 80826). One day later, media was replaced with media also containing 1 μg/ml doxycycline (Sigma, D9891) and 1x cumate (System Biosciences, QM100A-1), and sgRNA was transfected (see Supplementary Table 7). The day before imaging, media was replaced again with media containing 1 μg/ml doxycycline and 1x cumate. If the sgRNA was transduced (all data shown in Figs. 1-4), 260 μl of cell suspension [5 x 10^4^ cells/ml] were mixed with 40 μl of concentrated lentivirus, not titered, and plated into 8-well glass bottom chambers in media containing 8 μg/ml polybrene, 1 μg/ml doxycycline, and 1x cumate. Media was replaced 24 hours later with media containing 1 μg/ml doxycycline and 1x cumate and on the day of imaging, media was replaced ∼30 minutes before imaging with media containing 1 μg/ml doxycycline and 1x cumate. If the cell line had been previously transduced with sgRNA and selected for antibiotic resistance, 300 μl of cells were plated two days before imaging [5 x 10^4^ cells/ml] in 8-well glass bottom chambers and induced with 1 μg/ml doxycycline and 1x cumate, with media replacements each day. Cells in chambers were placed on the microscope in a live-cell incubator (Okolab, OKO-H301PI736ZR1/2S) and maintained at 37°C and 5% CO_2_ at 90% humidity.

Images were acquired with an exposure time of 50 ms at 20 Hz or 200 ms at 5 Hz, unless noted otherwise. Live cells were imaged on two different setups: (1) imaging with HILO and TIRF illumination was performed using an Olympus CellTIRF system with a Nikon 1.49 NA 100x oil objective, a 1.6x Optovar magnifier, and an Andor iXon Plus EMCCD camera, or (2) imaging with flat-field HILO illumination was performed on a custom-built microscope that was described in detail elsewhere^64^. Fixed cells were imaged using either a Zeiss LSM 800 confocal microscope or a Zeiss LSM 700 confocal microscope, both with a 1.40 NA 63x oil immersion objective. These cells were fixed for 10 min at room temperature with 4% formaldehyde solution made by diluting a 16% formaldehyde solution (Fisher Scientific, 50-980-487) in PBS, and then washed 1x and imaged in PBS.

### Dnase-seq data

Dnase-seq density data was downloaded from GEO Accession GSM4221655^82^, and the density was summed across the given genomic region, which was either a 100 kb window centered in the center of the repetitive locus, or only across the repetitive region itself.

### Tracking and MSD analysis

First, regions of interest of variable size in cells were manually selected to include a labeled locus (spot). The spot was localized with ThunderStorm^83^ using the following parameters: filter=[Wavelet filter (B-Spline)], scale=2.0, order=3, detector=[Local maximum], connectivity=8-neighbourhood, threshold=std(Wave.F1), estimator=[PSF: Integrated Gaussian], sigma=2.0, fitradius=3 method=[Weighted Least squares], and spurious localizations were filtered out by using the filtering parameters formula=[uncertainty_xy < 10 & offset > 0.00001]. Localizations were linked into a trajectory using trackpy.link^84^ using a maximum jump of 250 nm and frame gap of 3 frames, and trajectories shorter than 40 frames were eliminated from further analysis. Then, time-averaged (TA)-MSDs of individual trajectories were calculated using trackpy.ismd^84^, and the first 10 lags were fit to Eq. 1 describing fractional Brownian motion including localization uncertainty and motion blur^52,85^, by least-squares fitting in log-log space of D*, α, and localization precision, σ, with parameter bounds as follows: D* ∈ [0.000001, 1] µm^2^/s^α^, α ∈ [0.01, 2], σ ∈ [0.001, 0.1] µm. Fits with α < 0.05 or α > 1.95 indicated poor fits and were excluded from further analysis.

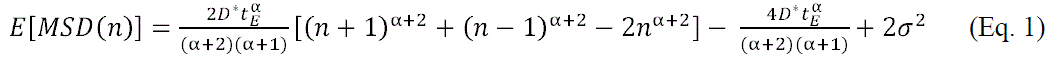

### Spot intensity measurements

To measure the spot intensities of labeled loci and single dSaCas9-ArrayG/Ns, the TrackMate plugin for FIJI was used^86^. For spot intensity histograms, the intensities of labeled loci and of single dSaCas9-ArrayG/Ns were normalized to the mean intensity of the single dSaCas9-ArrayG/N spots. For telomere intensity decay plots, spots were linked in TrackMate using a max distance of 5 pixels and a frame gap of 3 frames, and the intensity for each trajectory was normalized to the intensity of the first frame. In all directly compared experiments, the same laser intensity was used.

### Statistics and reproducibility

Tracking data were collected from 1-3 independent samples, each with its own lentiviral transduction and induction, and the data was pooled except for in Supplementary Fig. 4. Data that was directly compared (i.e. in Fig. 4) were acquired from individual samples transduced with the same lentiviral preparation and plated, induced, and imaged on the same days. Information on the number of trajectories and cells for each experiment can be found in each figure caption. For Fig. 1f, data shows mean ± std. For Fig. 2e, data shows mean ± SEM. In Figs. 1c,g, 2d, and 3e, data shows violin plots (representing the kernel density estimation (KDE)) overlaid with the quartiles and whiskers of a box plot. In Figs. 4a,b,e, data shows split violin plots (representing the KDE) where the dashed lines indicate the quartiles. All p-values are the results of two-sided Mann-Whitney U tests. No statistical method was used to predetermine sample size throughout this work.

## Data availability

Localizations and trajectories generated and analyzed during the current study and source data that can be used for generating the graphs in this work are provided with this paper and on GitHub [https://github.com/Gustavsson-Lab/chromtrack/tree/main].

## Code availability

The sgRNA were selected using the design tool CHOPCHOP^81^ [https://chopchop.cbu.uib.no]. Loci were localized using the open-source FIJI plugin ThunderSTORM^83^ [https://github.com/zitmen/thunderstorm/releases/tag/v1.3/]. Localizations were linked into a trajectory using trackpy.link^84^ [https://soft-matter.github.io/trackpy/dev/generated/trackpy.link.html]. MSDs were calculated using trackpy.imsd^84^ [https://soft-matter.github.io/trackpy/dev/generated/trackpy.motion.imsd.html#trackpy.motion.imsd]. Spot intensities were analyzed using the open-source FIJI plugin TrackMate^86^ [https://github.com/trackmate-sc/TrackMate]. Fits were performed and plots were generated using custom code in Python with common packages, available on GitHub [https://github.com/Gustavsson-Lab/chromtrack/tree/main].

## Acknowledgements

This work was supported by partial financial support from the National Institute of General Medical Sciences of the National Institutes of Health grant R35GM155365, the National Science Foundation grant 2441757, and startup funds from the Cancer Prevention and Research Institute of Texas grant RR200025 to A.-K.G. J.T.L. acknowledges support from the Nucleome Initiative, NCI Physical Sciences-Oncology Center, grant U54CA143836. A.R. acknowledges partial financial support from the Bioelectronics NSF Research Traineeship (NRT) Program. This work was conducted in part using resources and equipment available through the Shared Equipment Authority at Rice University. We thank Dr. Alloysius Budi Utama at the Shared Equipment Authority for his input and guidance. We thank Dr. Robert Singer for providing U2OS p21-MS2 MCP-EGFP cells.

## Author contributions

R.P.G. conceived the idea. R.P.G., A.R., and A.-K.G. designed experiments. R.P.G., A.R., and M.K.N. created plasmids and cell lines. R.P.G., A.R., Q.S., and L.Y. performed imaging experiments. A.R. analyzed data. I.B.H., J.T.L., and A.-K.G. supervised the research. R.P.G., A.R., and A.-K.G. wrote the paper.

