## Supplementary Material for "Robust fluorescent labeling and tracking of endogenous non-repetitive genomic loci"

[1] *Systems, Synthetic, and Physical Biology PhD Program, Rice University, Houston, TX, USA*
[2] *Department of Chemistry, Rice University, Houston, TX, USA*
[3] *Howard Hughes Medical Institute, University of California, Berkeley, Berkeley, CA, USA*
[4] *Department of Bioengineering, Stanford University, Stanford, CA, USA*
[5] *Center for Theoretical Biological Physics, Rice University, Houston TX, USA*
[6] *Department of Bioengineering, Rice University, Houston, TX, USA*
[7] *Department of BioSciences, Rice University, Houston, TX, USA*
[8] *Department of Electrical and Computer Engineering, Rice University, Houston, TX, USA*
[9] *Smalley Curl Institute, Rice University, Houston, TX, USA*
[10] *Center for Nanoscale Imaging Sciences, Rice University, Houston, TX, USA*
[11] *Department of Cancer Biology, University of Texas MD Anderson Cancer Center, Houston, TX, USA*
[*] *These authors contributed equally.*

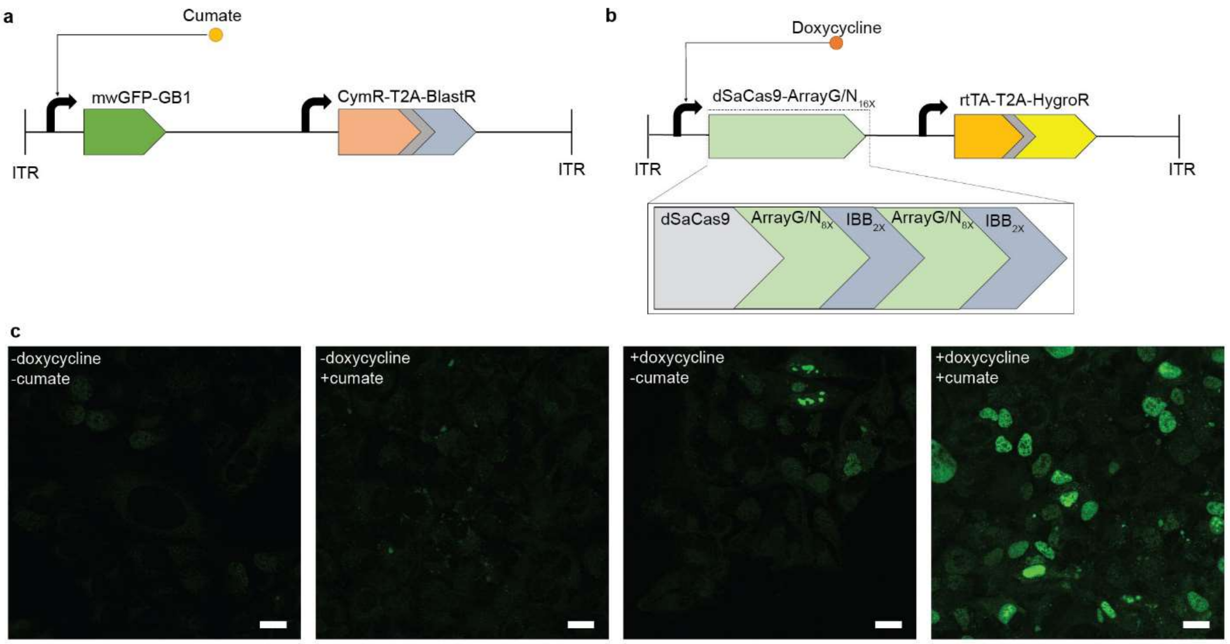


**Supplementary Figure 1: Overview of dSaCas9-ArrayG/N. a** Construct schematic of mwtGFP-GB1 driven by a cumate-inducible promoter. The construct also contains CymR which enables cumate induction and a blasticidin resistance marker, all sandwiched between piggyBac transpose inverted terminal repeats (ITRs). **b** Construct schematic of dSaCas9-ArrayG/$N_{16X}$ driven by doxycycline-inducible promoter. The construct also contains rtTA3 which enables doxycycline induction and a hygromycin resistance marker, all sandwiched between ITRs. **c** Representative images of fixed U2OS cells stably integrated with both constructs without induction, with only cumate induction, with only doxycycline induction, and with both cumate and doxycycline induction, respectively from left to right. The images were acquired on the same day with the same acquisition settings. All images are scaled such that their displayed range matches the minimum (0) and maximum intensity values of the leftmost image. Scale bars are 25 μm.

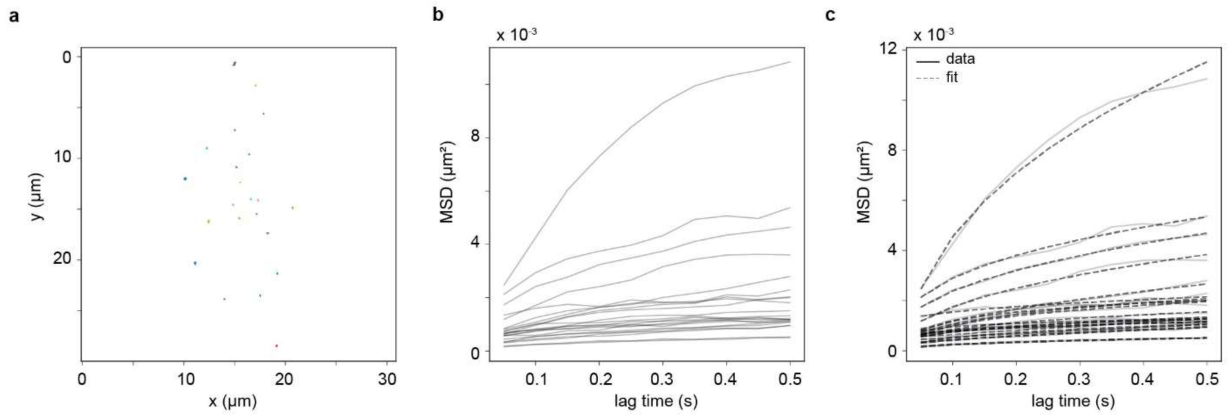


**Supplementary Figure 2: Example mean squared displacement (MSD) analysis. a** Trajectories from a 20 Hz telomere tracking experiment. **b** MSDs of individual trajectories. **c** Same as **b** but overlaid with the fits of each individual MSD (dashed lines).

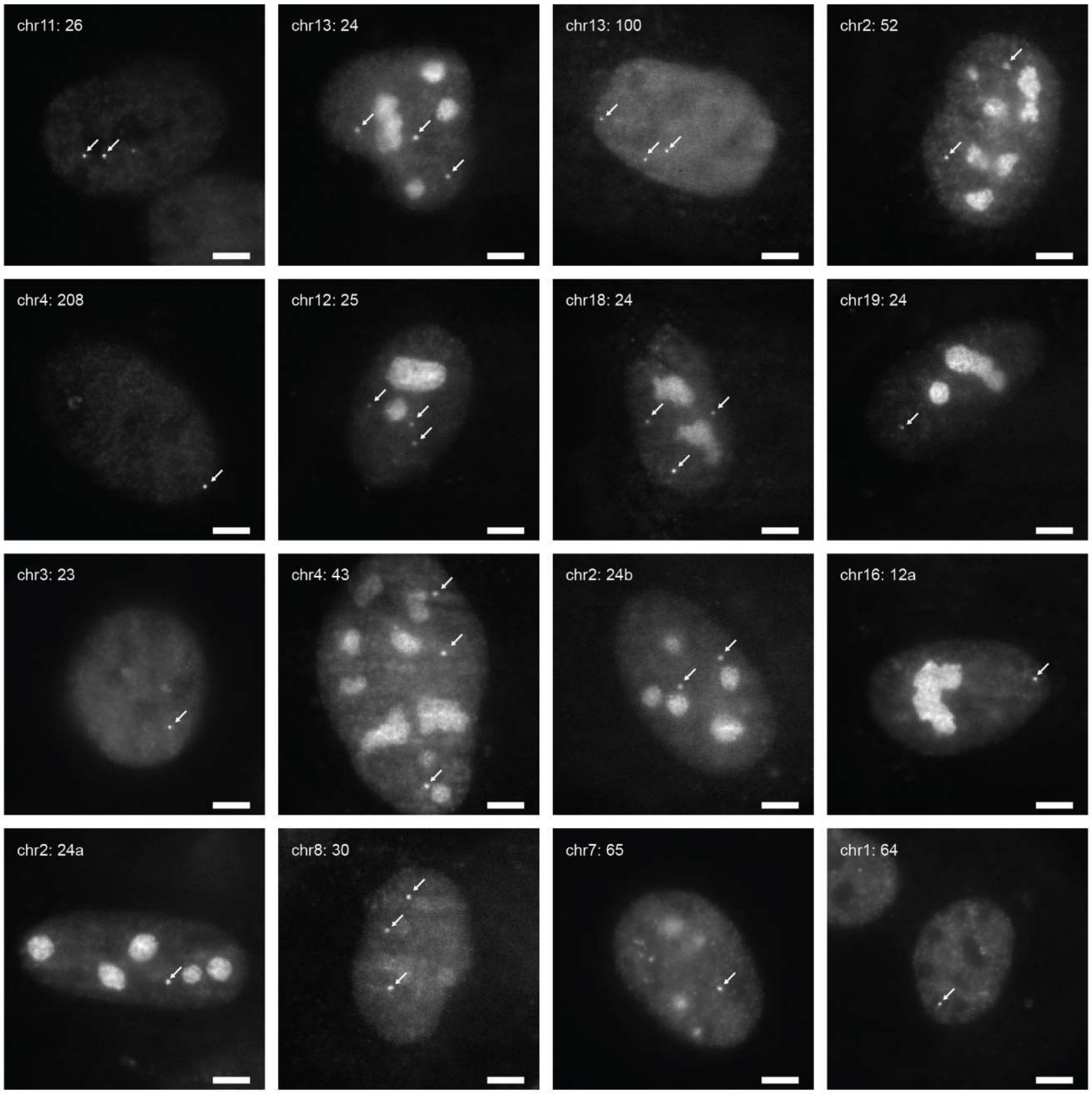


**Supplementary Figure 3: Imaging repetitive loci with dSaCas9-ArrayG/N.** Representative images of cells labeled for the indicated (by chromosome and repeat number) repetitive locus. All cells are labeled by dSaCas9-ArrayG/$N_{8X}$ except chr3: 23, chr7: 65, and chr1: 64, which are labeled by dSaCas9-ArrayG/$N_{16X}$. White arrows point to the loci. Intensity normalized independently for each image. Scale bars are 5 μm.

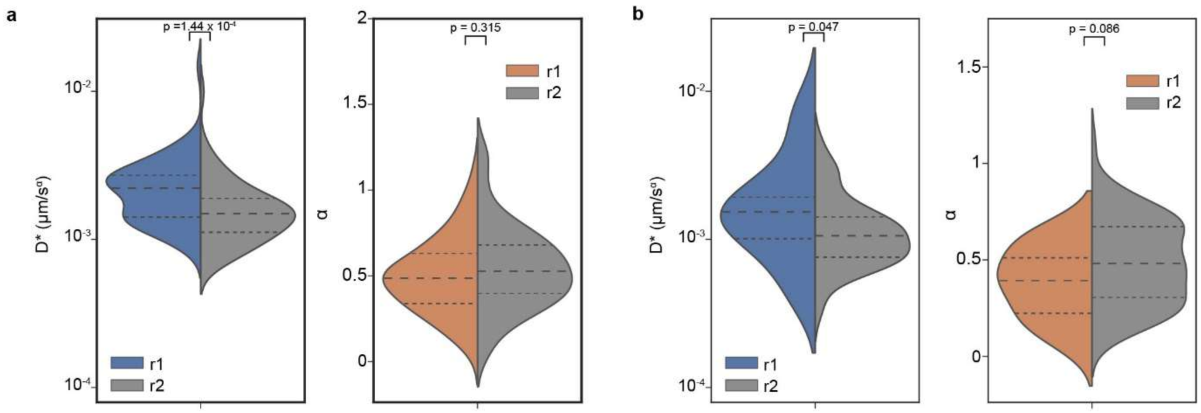


**Supplementary Figure 4: Analysis of replicate experiments. a** Mean square displacement analysis of repeat locus chr13: 24 tracked at 5 Hz, shown as split violin plots of the (*left*) effective diffusion coefficient, D*, and (*right*) anomalous exponent, α, split between experiments (r1 and r2) performed on two different days ($n$ = 67 trajectories for r1 and 66 trajectories for r2). **b** Mean square displacement analysis of the non-repetitive locus *PCNA* tracked at 5 Hz, shown as split violin plots of the (*left*) effective diffusion coefficient and (*right*) anomalous exponent, split between experiments performed on two different days ($n$ = 16 trajectories for r1 and 93 trajectories for r2). All p-values are the result of two-sided Mann-Whitney U tests.

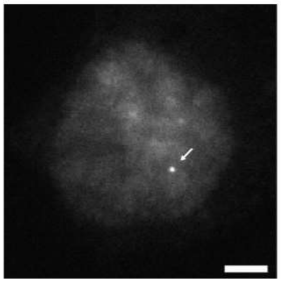

**Supplementary Figure 5: Proof of concept of non-repetitive imaging with 6 sgRNAs.** Representative image of *VCL* labeled with 6 sgRNAs and dSaCas9-ArrayG/$N_{16X}$ expressed from a tandem U6 construct. Scale bar is 5 μm.

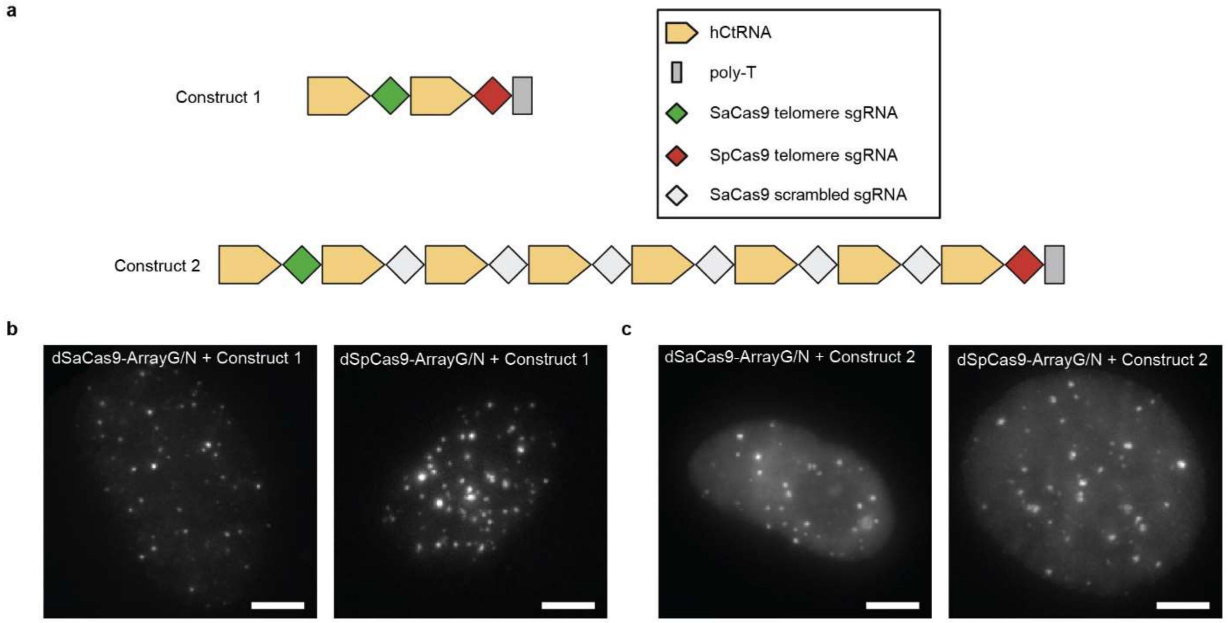


**Supplementary Fig 6: Drive-and-process array produces sufficient sgRNA for imaging. a** Construct diagrams of the two constructs used for validation. In Construct 1, the array expresses a telomere-targeting sgRNA for SaCas9 and a telomere-targeting sgRNA for SpCas9. Construct 2 is the same as Construct 1, except that 6 scrambled SaCas9 sgRNAs have been placed between the two telomere sgRNAs. **b** Representative images of cells transfected with Construct 1 and expressing (*left*) dSaCas9-ArrayG/$N_{16X}$ or (*right*) dSpCas9-ArrayG/$N_{16X}$. **c** Representative images of cells transfected with Construct 2 and expressing (*left*) dSaCas9-ArrayG/$N_{16X}$ or (*right*) dSpCas9-ArrayG/$N_{16X}$. Images are shown as maximum intensity projections of z-stacks covering the full nucleus. Scale bars are 5 μm.

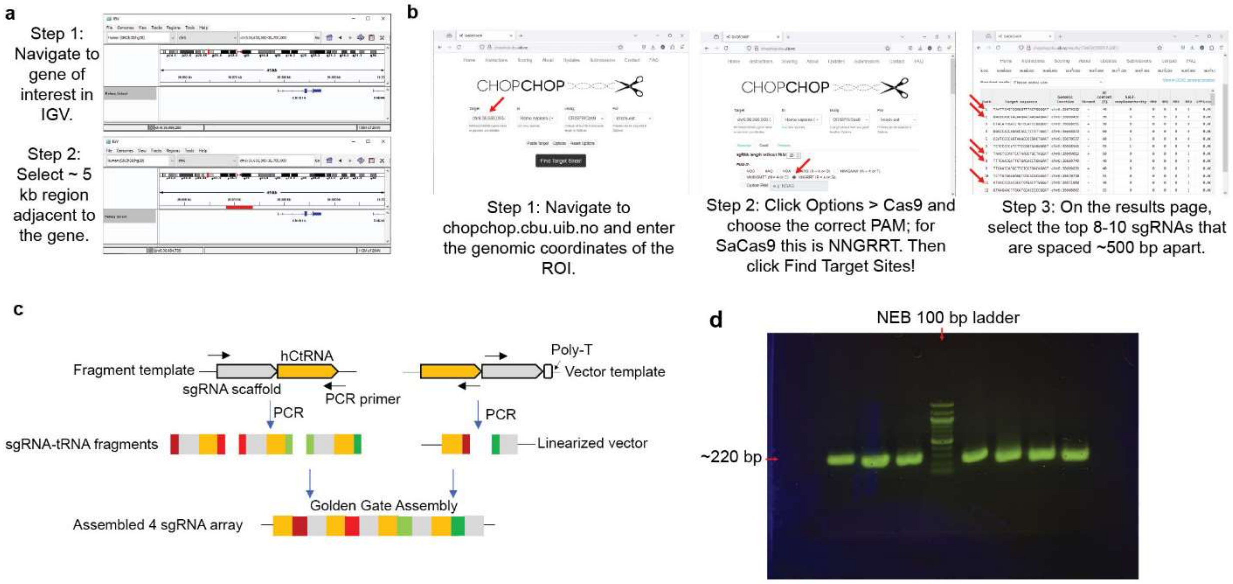


**Supplementary Figure 7: Process of cloning a drive-and-process array. a** Region selection in the Integrated Genome Viewer (IGV). **b** sgRNA design in CHOPCHOP[1]. In step 3, only the first 5 sgRNAs are pointed to with red arrows. **c** Cartoon schematic of the Golden Gate Assembly (GGA) process, modified from Yuan and Gao[2]. Primers are designed to amplify n-1 fragment templates and 1 vector template for an n-sgRNA array, such that the amplicons contain ~ half of each sgRNA (indicated by colors in the amplicons) and BsaI cut sites to generate compatible overhangs for GGA. GGA then assembles the n-sgRNA array. **d** A representative image of an agarose gel of the n-1 fragment PCR amplicons when cloning an 8-sgRNA array.

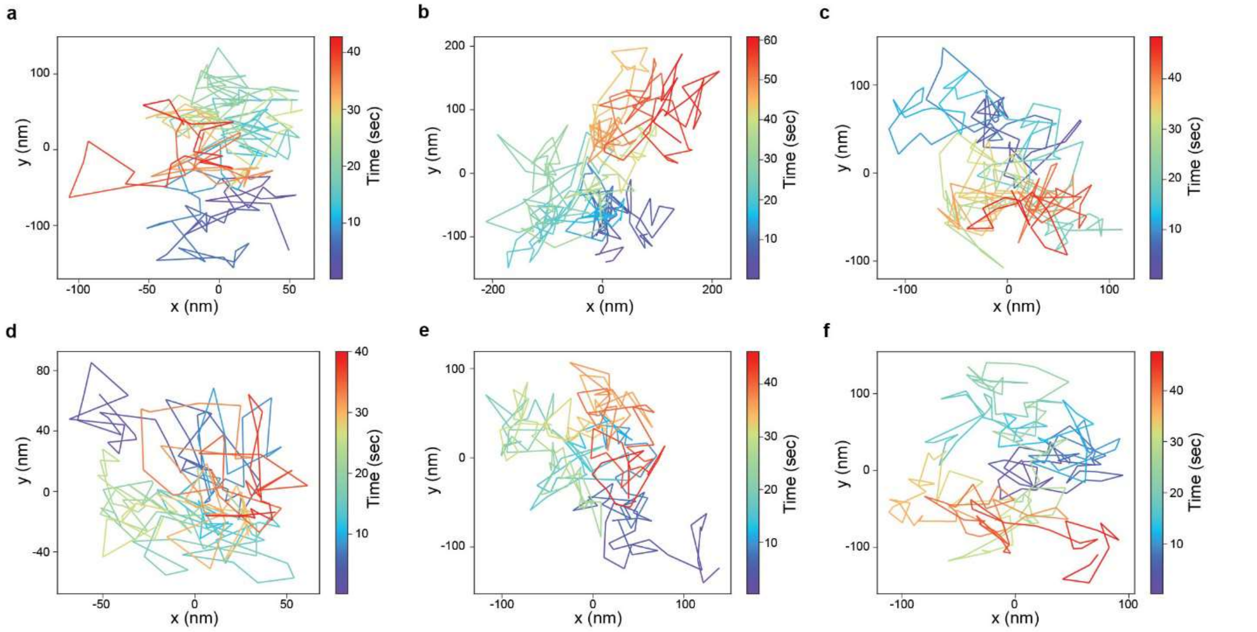

**Supplementary Figure 8: dSaCas9-ArrayG/N enables many-frame trajectories of non-repetitive loci.** Representative 5 Hz trajectories, colored by time, for non-repetitive loci **a** *CCNB1*, **b** *CDKN1A*, **c** *PCNA*, **d** *RUNX2*, **e** *TLN1*, and **f** *VCL*.

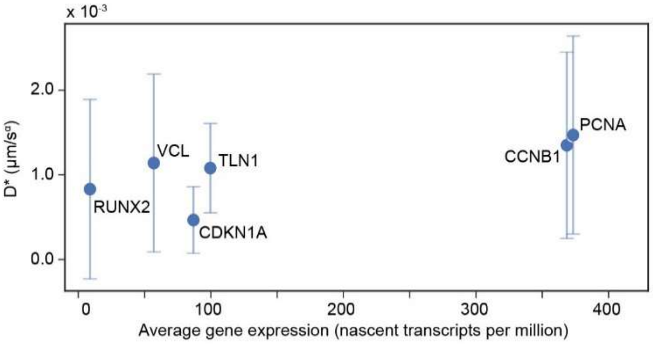


**Supplementary Figure 9: Gene expression and diffusion.** Scatterplot of the effective diffusion coefficient of each gene tracked at 5 Hz (as in Fig. 3) against the average gene expression in U2OS cells (sourced from the Human Protein Atlas) (data shown as mean ± standard deviation, $n$ = 39, 26, 111, 23, 41, 81 trajectories across 30, 20, 78, 21, 34, 43 cells for *CCNB1*, *CDKN1A*, *PCNA*, *RUNX2*, *TLN1*, and *VCL*, respectively).

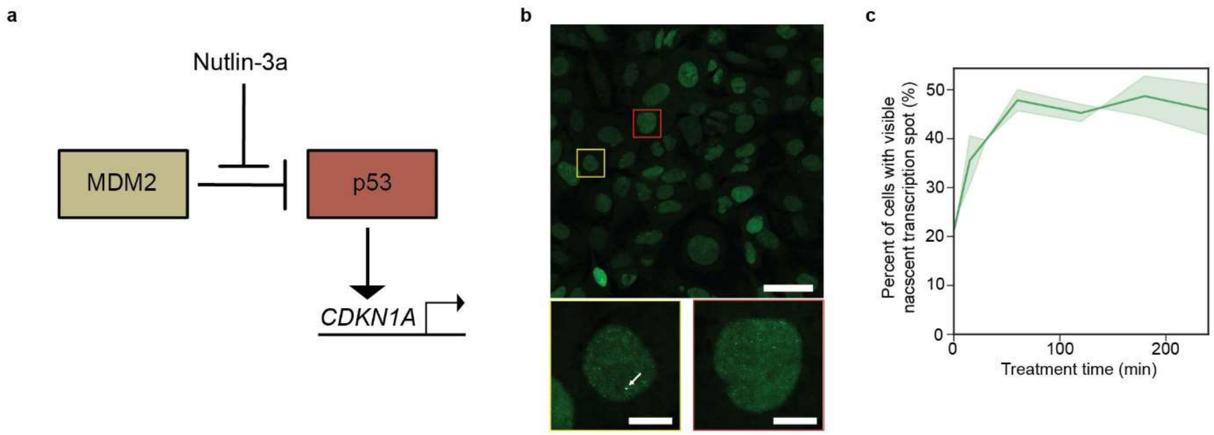


**Supplementary Figure 10: Validation of *CDKN1A* activation by Nutlin-3a. a** Schematic of Nutlin-3a mediated activation of *CDKN1A* transcription. Nutlin-3a interferes with p53 inhibition by MDM2, enabling p53 to transcriptionally activate target genes including *CDKN1A*[3]. **b** Representative stitched fields of view of fixed U2OS p21-MS2 MCP-EGFP cells after 15 min treatment with 10 µM Nutlin-3a, shown as a max-intensity projection of a z-stack covering the full nuclei. Scale bar is 25 µm. Bounded boxes are zoom ins on the selected cells (scale bars are 5 µm); yellow is a cell with a visible NTS (white arrow points to NTS); red is a cell without a visible spot. **c** Quantification of the fraction of cells with a visible NTS for different treatment times (data shown as the mean of two independent experiments, where the shaded bounds are the separate experiments. *n* = 21, 26, 31, 48, 57, 65, 71, 69 transcribing cells out of 99, 100, 91, 121, 118, 144, 146, and 147 cells for time points 0, 5, 15, 30, 60, 120, 180, and 240 minutes, respectively).

**Supplementary Table 1: Telomere dynamics at 20 Hz.** The fitted effective diffusion coefficient (D*), anomalous exponent (α), and localization uncertainty (σ) of telomeres labeled by the indicated dCas9 tags and tracked at 20 Hz are shown as the mean ± standard deviation of the indicated number of trajectories.

| dCas9 Label | *n* (trajectories) | D* ($\mu m^2/s^a$) | α | σ (nm) |
|---|---|---|---|---|
| dSaCas9-ArrayG/$N_{16X}$ | 1276 | $1.53 \pm 1.50 \times 10^{-3}$ | 0.62 ± 0.31 | 12.6 ± 4.3 |
| dSaCas9-ArrayG/$N_{8X}$ | 884 | $2.17 \pm 3.04 \times 10^{-3}$ | 0.63 ± 0.30 | 13.8 ± 5.7 |
| dSpCas9-ArrayG/$N_{16X}$ | 265 | $2.24 \pm 2.76 \times 10^{-3}$ | 0.65 ± 0.34 | 11.4 ± 5.5 |
| dSpCas9-ArrayG/$N_{8X}$ | 294 | $2.21 \pm \pm 1.96 \times 10^{-3}$ | 0.69 ± 0.34 | 13.6 ± 5.2 |
| dSpCas9-EGFP | 34 | $1.57 \pm 1.14 \times 10^{-3}$ | 0.67 ± 0.42 | 15.2 ± 7.6 |

**Supplementary Table 2: Locally repetitive dSaCas9 targets.** The fitted effective diffusion coefficient (D*), anomalous exponent (α), and localization uncertainty (σ) of the indicated repetitive loci and tracked at 5 Hz are shown as the mean ± standard deviation of the indicated number of trajectories. Also included are the locus location (Chromosome and Center Position in hg38), size, DNAse score in a 100 kb window around the center of the locus and within the repeat region, and the spacer sequence used in the sgRNA for labeling.

| Chromosome | Copy # | Center Position (hg38) | Size (bp) | DNAse in 100 kb window | DNAse in repeat region | Spacer Sequence | *n* (tracks) | D* ($\mu m^2/s^a$) x $10^3$ | α | σ (nm) |
|---|---|---|---|---|---|---|---|---|---|---|
| 4 | 208 | 186966452.5 | 23931 | 4.850994061 | 0.029106077 | atggaagaggcaggcgacat | 32 | 3.73 ± 3.3 | 0.57 ± 0.32 | 20.5 ± 6.7 |
| 13 | 100 | 112298054.5 | 50177 | 4.322188208 | 0.015226349 | cttacctggctccatcctga | 41 | 5.76 ± 6.2 | 0.56 ± 0.20 | 20.2 ± 8.4 |
| 1 | 51 | 201260979.5 | 12663 | 7.314115872 | 0.068786772 | gatggagtgagatggagtga | 127 | 4.50 ± 3.8 | 0. 42 ± 0.20 | 7.7 ± 8.0 |
| 2 | 24 | 17963962 | 11284 | 5.763776375 | 0.050993866 | atccaggagagatgactgca | 63 | 1.31 ± 1.1 | 0.56 ± 0.23 | 10.1 ± 6.0 |
| 16 | 12 | 81867163 | 11024 | 6.001550484 | 0.052821234 | gctggcctctcccttgctcc | 88 | 1.39 ± 1.0 | 0.61 ± 0.30 | 9.6 ± 7.2 |
| 16 | 12 | 89014968.5 | 10871 | 6.333642495 | 0.045012052 | atagatgggtggatgagtgc | 18 | 1.04 ± 1.0 | 0.57 ± 0.25 | 9.9 ± 3.8 |
| 2 | 52 | 1107531.5 | 13015 | 2.468963126 | 0.026153514 | cacgtgctgtcactcggtg | 50 | 1.74 ± 1.1 | 0.54 ± 0.25 | 9.0 ± 5.6 |
| 18 | 24 | 9819946.5 | 10805 | 12.81194434 | 0.139206286 | gtagatgctgtcctccatcc | 143 | 1.97 ± 1.6 | 0.48 ± 0.21 | 7.3 ± 6.0 |
| 13 | 24 | 29558619.5 | 11795 | 3.861416683 | 0.035147773 | gggagtgaatgtgagtgagg | 67 | 2.38 ± 1.8 | 0.50 ± 0.22 | 8.9 ± 6.4 |
| 2 | 24 | 92111198.5 | 36741 | 0.56351002 | 0 | tgatgtgtgtactcaactaa | 38 | 1.57 ± 0.9 | 0.51 ± 0.21 | 8.5 ± 4.7 |
| 4 | 43 | 49130370.5 | 54065 | 6.07559722 | 0.077670303 | tggaacggaacggaacggaa | 14 | 1.21 ± 0.7 | 0.47 ± 0.21 | 9.3 ± 4.9 |
| 16 | 29 | 88955144.5 | 11653 | 6.705593154 | 0.054507999 | gtggggtcaggacctttggg | 84 | 1.44 ± 0.6 | 0.56 ± 0.25 | 8.4 ± 5.8 |
| 12 | 25 | 124467968 | 12024 | 11.53202007 | 0.047889733 | atgatgggggtgatgaggat | 75 | 1.87 ± 1.1 | 0.47 ± 0.25 | 7.7 ± 6.1 |
| 11 | 26 | 65971196.5 | 10881 | 8.889709572 | 0.063691586 | tgagctgaggaggggatgg | 21 | 4.92 ± 4.7 | 0.34 ± 0.15 | 6.6 ± 6.8 |
| 19 | 24 | 2713752 | 11320 | 3.588938805 | 0.022958964 | gatgaagggtggaggaggtg | 51 | 1.65 ± 1.1 | 0.54 ± 0.31 | 9.3 ± 5.9 |
| 3 | 30 | 77292817.5 | 17069 | 3.703874185 | 0.030227002 | aaagtaaaattgacggttaa | 102 | 1.44 ± 0.7 | 0.54 ± 0.21 | 7.9 ± 5.1 |
| 8 | 30 | 1399660 | 22174 | 1.42337782 | 0.00670808 | gcactcaatacacgactcag | 21 | 1.17 ± 0.9 | 0.52 ± 0.24 | 6.3 ± 4.5 |
| 3 | 23 | 196058291 | 12688 | 4.225979584 | 0.038536444 | gaagtgtcggtgacaggaag | 26 | 1.86 ± 1.8 | 0.43 ± 0.19 | 3.8 ± 4.8 |
| 1 | 64 | 1077639 | 3572 | 6.880488062 | 0.229613018 | gcctcagcctcccccca | 14 | 0.48 ± 0.53 | 0.61 ± 0.23 | 6.9 ± 5.4 |
| 7 | 65 | 158336302.5 | 12667 | 2.937123072 | 0.0108103 | gctcttatggtgagagt | 19 | 0.37 ± 0.35 | 0.57 ± 0.22 | 4.8 ± 3.2 |

**Supplementary Table 3: Non-repetitive sgRNAs details.** The spacer sequences used for sgRNAs in non-repetitive labeling are shown above the starting position of the target site in genomic coordinates (hg38).

| ***CCNB1-3+5'*** | ***CDKN1A-5'*** | ***PCNA-3'*** | ***RUNX2-5'*** | ***TLN1-3+5'*** | ***VCL-3'*** |
|---|---|---|---|---|---|
| cgaggtttcaccatgttagc<br>chr5:69,164,745 | gacgcagccacaagaataag<br>chr6:36,668,497 | ctgcactccagcctggtcaa<br>chr20:5,114,135 | cccccaagtcacacataggt<br>chr6:45,403,753 | gtgactaggccttgtacatc<br>chr9:35,730,934 | acaatgagccttgcccattg<br>chr10:74121704 |
| tcccagtgctttgtgaggcc<br>chr5:69164057 | taagtccagtcctaagctgc<br>chr6:36,669,013 | gagttctagttactggacat<br>chr20:5,113,890 | tagctgaagctagtataaat<br>chr6:45,402,211 | atgttggcatggtcggtgtt<br>chr9:35,731,526 | gctcctggcgtacattctac<br>chr10:74,121,909 |
| ggcgggcaaatcacaaggtc<br>chr5:69,163,773 | tttgaaggcagctcacagga<br>chr6:36,669,684 | gctattcaagggaagtatag<br>chr20:5,113,580 | aggagaaccattggcagtgg<br>chr6:45,400,299 | gcctctcttgcgtgtccggt<br>chr9:35,732,002 | ggtagccaaacttaggcaaa<br>chr10:74,122,090 |
| caggagtatcatttgagccc<br>chr5:69,163,211 | tccttctcctatcaatgggt<br>chr6:36,670,079 | gcttcccagggaggcagaag<br>chr20:5,113,397 | gccatacggtcagataggct<br>chr6:45,404,836 | tgtcccaaatggagctggaa<br>chr9:35,732,765 | aagaagaatggagccacaaa<br>chr10:74,122,355 |
| ctcagggattgcttgagctt<br>chr5:69,162,917 | taattcagtcggggtttact<br>chr6:36,670,535 | tcctgcaggtgacactcggt<br>chr20:5,113,187 | gtggttagtagaagataaca<br>chr6:45,401,227 | taacgctgctgtacctcgat<br>chr9:35,697,540 | tccaaaaccacggtgatcgg<br>chr10:74,122,559 |
| tgcatgcccttcaataacgt<br>chr5:69,180,458 | gagtgccccaccaacttcta<br>chr6:36,671,101 | ctccctggcaaggtctgaag<br>chr20:5,112,766 | tcggtaaagctcaccacaca<br>chr6:45,400,751 | tctcaggcacaacttcccca<br>chr9:35,697,147 | taaccatggcttaactggtg<br>chr10:74,122,886 |
| gggagagtcacttgaggcc<br>chr5:69,180,870 | ataacccaacacatgtgact<br>chr6:36,671,860 | tcgtaacattcaggtgctga<br>chr20:5,112,666 | tttatctttagcagttggcc<br>chr6:45,403,166 | cttttagcgctaccctaggc<br>chr9:35,696,798 | tctcgggacagatactggtt<br>chr10:74,123,373 |
| tctagctctgttgcccaagc<br>chr5:69,181,008 | gagagctaggcccttacacc<br>chr6:36,672,263 | tcgggtccctaaactctagc<br>chr20:5,111,892 | ggagacatcctagagcagga<br>chr6:45,404,374 | caagctttccctaatgtggt<br>chr9:35,696,456 | agtgataacctctctcctaa<br>chr10:74,123,506 |
| gccactgtgcccagcccttc<br>chr5:69,181,266 | | gattcttctgcttcagcctc<br>chr20:5,111,504 | | agctgtcctgggctcctagc<br>chr9:35,695,981 | ctattaggtaccaagtgagt<br>chr10:74,123,751 |
| tccacagtttgagcacagca<br>chr5:69,181,523 | | | | gagaagacagggtgtagaaa<br>chr9:35,729,974 | ttaagattcccacacaggca<br>chr10:74,124,081 |

**Supplementary Table 4: Gene dynamics at 5 Hz.** The fitted effective diffusion coefficient (D*), anomalous exponent (α), and localization uncertainty (σ) of the indicated genes labeled by the indicated dCas9 tags and tracked at 5 Hz are shown as the mean ± standard deviation of the indicated number of trajectories.

| Gene | dCas9 Label | *n* (trajectories) | D* ($\mu m^2/s^a$) | α | σ (nm) |
|---|---|---|---|---|---|
| *CCNB1* | dSaCas9-ArrayG/$N_{8X}$ | 39 | $1.37 \pm 1.14 \times 10^{-3}$ | 0.53 ± 0.27 | 7.5 ± 5.4 |
| *CDKN1A* | dSaCas9-ArrayG/$N_{16X}$ | 26 | $0.478 \pm 0.39 \times 10^{-3}$ | 0.55 ± 0.21 | 4.7 ± 3.1 |
| *PCNA* | dSpCas9-ArrayG/$N_{8X}$ | 11 | $1.43 \pm 1.15 \times 10^{-3}$ | 0.47 ± 0.22 | 7.2 ± 5.3 |
| *RUNX2* | dSpCas9-ArrayG/$N_{16X}$ | 23 | $0.849 \pm 1.1 \times 10^{-3}$ | 0.50 ± 0.23 | 6.0 ± 9.7 |
| *TLN1* | dSpCas9-ArrayG/$N_{8X}$ | 41 | $1.1 \pm 0.52 \times 10^{-3}$ | 0.54 ± 0.25 | 7.7 ± 5.0 |
| *VCL* | dSpCas9-ArrayG/$N_{8X}$ | 81 | $1.05 \pm 0.82 \times 10^{-3}$ | 0.53 ± 0.18 | 4.5 ± 3.2 |

**Supplementary Table 5: Transcription-independent gene dynamics at 20 Hz.** The fitted effective diffusion coefficient (D*), anomalous exponent (α), and localization uncertainty (σ) of the indicated genes in the indicated transcriptional condition and tracked at 20 Hz are shown as the mean ± standard deviation of the indicated number of trajectories.

| Gene | Condition | *n* (trajectories) | D* ($\mu m^2/s^a$) x $10^3$ | α | σ (nm) |
|---|---|---|---|---|---|
| *CDKN1A* | Control | 22 | 1.09 ± 0.97 | 0.42 ± 0.27 | 8.4 ± 7.6 |
| *CDKN1A* | Triptolide | 18 | 0.89 ± 0.42 | 0.41 ± 0.16 | 9.6± 4.6 |
| *CDKN1A* | Control | 15 | 1.34 ± 1.30 | 0.43 ± 0.26 | 14.6 ± 8.0 |
| *CDKN1A* | Nutlin-3a | 23 | 2.21± 2.90 | 0.35 ± 0.16 | 10.0 ± 7.7 |
| *PCNA* | Control | 31 | 1.72 ± 1.09 | 0.43 ± 0.20 | 13.7 ± 6.1 |
| *PCNA* | Triptolide | 69 | 3.30 ± 2.57 | 0.36 ± 0.18 | 13.9 ± 7.5 |
| *VCL* | Control | 35 | 3.20 ± 2.63 | 0.43 ± 0.21 | 17.6 ± 9.6 |
| *VCL* | Triptolide | 50 | 3.96 ± 3.43 | 0.53 ± 0.31 | 15.3 ± 7.8 |
| *VCL* | Other | 40 | 1.63 ± 1.31 | 0.47 ± 0.19 | 15.8 ± 6.8 |
| *VCL* | Overlap | 20 | 1.98 ± 2.34 | 0.44 ± 0.26 | 13.7 ± 8.1 |

### Supplementary Table 6: List of plasmids used in this study.

| Code | Full Name | Description | Source | Availability |
|---|---|---|---|---|
| pRPG001 | pbTRE3G_dSaCas9-ArrayG/$N_{16X}$ | Dox-inducible expression of dSaCas9-ArrayG/$N_{16X}$ | This study. | Will be deposited to Addgene. |
| pRPG001.5 | pbTRE3G_dSaCas9-ArrayG/$N_{8X}$ | Dox-inducible expression of dSaCas9-ArrayG/$N_{8X}$ | This study. | Will be deposited to Addgene. |
| pRPG002 | pbTRE3G_dSpCas9-ArrayG/$N_{16X}$ | Dox-inducible expression of dSpCas9-ArrayG/$N_{16X}$ | Gustavsson, A.-K., Ghosh, R. P., Petrov, P. N., Liphardt, J. T. & Moerner, W. E. Fast and parallel nanoscale three-dimensional tracking of heterogeneous mammalian chromatin dynamics. Mol. Biol. Cell 33, 1–11 (2022). | Will be deposited to Addgene. |
| pRPG007 | pbQM_mwGFP-GB1 | Cumate-inducible expression of mwGFP-GB1 | Gustavsson, A.-K., Ghosh, R. P., Petrov, P. N., Liphardt, J. T. & Moerner, W. E. Fast and parallel nanoscale three-dimensional tracking of heterogeneous mammalian chromatin dynamics. Mol. Biol. Cell 33, 1–11 (2022). | Will be deposited to Addgene. |
| pAR101 | hCtRNA_VT-dSp | Vector template for optimized dSpCas9 DAP sgRNA array | This study. | Will be deposited to Addgene. |
| pAR102 | hCtRNA_FT-dSp | Fragment template for optimized dSpCas9 DAP sgRNA array | This study. | Will be deposited to Addgene. |
| pAR103 | hCtRNA_VT-dSa | Vector template for optimized dSaCas9 DAP sgRNA array | This study. | Will be deposited to Addgene. |
| pAR104 | hCtRNA_FT-dSa | Fragment template for optimized dSaCas9 DAP sgRNA array | This study. | Will be deposited to Addgene. |
| pAR105 | pDAP_SaTelo-SpTelo | DAP array expressing Sa- and SpCas9 telomere sgRNA | This study. | Available upon request. |
| pAR112 | pDAP_SaTelo-6xSaRandom-SpTelo | DAP array expressing SaCas9 telomere sgRNA, 6 nontargeting SaCas9 sgRNAs, and SpCas9 telomere sgRNA | This study. | Available upon request. |
| pAR111 | pDAP_Sa-CDKN1A-5UTR | DAP array expressing 8 sgRNAs for labeling upstream of CDKN1A | This study. | Available upon request. |
| pAR109 | pDAP_Sa-RUNX2-5UTR | DAP array expressing 8 sgRNAs for labeling upstream of RUNX2 | This study. | Available upon request. |
| pAR211 | pLP_DAP_Sa-CDKN1A-5UTR | Lentiviral transfer vector of DAP array expressing 8 sgRNAs for labeling upstream of CDKN1A | This study. | Available upon request. |
| pAR206 | pLP_DAP_Sa-RUNX2-5UTR | Lentiviral transfer vector of DAP array expressing 8 sgRNAs for labeling upstream of RUNX2 | This study. | Available upon request. |
| pAR220 | pLP_sasgRNA | Lentiviral transfer vector template for cloning in U6-driven SaCas9 sgRNAs using BbsI golden gate assembly. Used for repeat imaging (Fig 2). | This study. | Will be deposited to Addgene. Plasmids with the sgRNA spacer cloned in and used for imaging in Figures 1 and 2 are available upon request. |
| pRPG100 | pLTUB_sasgRNA-tRNA | Lentiviral transfer vector for cloning of PTG arrays. | This study. | Available upon request. |
| pRPG101 | pLTUB_10xPTG-CCNB1 | Lentiviral transfer vector of PTG array expressing 10 sgRNAs for labeling flanking CCNB1. | This study. | Available upon request. |
| pRPG102 | pLTUB_10xPTG-PCNA | Lentiviral transfer vector of PTG array expressing 10 sgRNAs for labeling downstream of PCNA. | This study. | Available upon request. |
| pRPG103 | pLTUB_10xPTG-TLN1 | Lentiviral transfer vector of PTG array expressing 10 sgRNAs for labeling flanking TLN1. | This study. | Available upon request. |
| pRPG104 | pLTUB_10xPTG-VCL | Lentiviral transfer vector of PTG array expressing 10 sgRNAs for labeling downstream of VCL. | This study. | Available upon request. |

| | | | | |
|---|---|---|---|---|
| hCtRNA_VT | hCtRNA_VT | Vector template for dSpCas9 DAP sgRNA array | Yuan, Q., Gao, X. Multiplex base- and prime-editing with drive-and-process CRISPR arrays. Nat Commun 13, 2771 (2022). https://doi.org/10.1038/s41467-022-30514-1 | Addgene #186716 |
| hCtRNA_FT | hCtRNA_FT | Fragment template for dSpCas9 DAP sgRNA array | Yuan, Q., Gao, X. Multiplex base- and prime-editing with drive-and-process CRISPR arrays. *Nat Commun* 13, 2771 (2022). https://doi.org/10.1038/s41467-022-30514-1 | Addgene #186715 |
| pLH-spsgRNA2 | pLH-spsgRNA2 | Lentiviral transfer vector template for cloning in U6-driven SpCas9 sgRNAs using BbsI golden gate assembly | Ma et al Proc Natl Acad Sci U S A. 2015 Mar 10;112(10):3002-7. doi: 10.1073/pnas.1420024112. Epub 2015 Feb 23. | Addgene #64114 |
| pMDLG/pRRe | pMDLG/pRRE | 3rd gen lentiviral packaging plasmid | Dull et al J Virol. 1998 Nov . 72(11):8463-71. | Addgene #12251 |
| pRSV-Rev | pRSV-Rev | 3rd gen lentiviral paclaging plasmid | Dull et al J Virol. 1998 Nov . 72(11):8463-71. | Addgene #12253 |
| pMD2.G | pMD2.G | VSV-G envelope expressing plasmid, for lentiviral production | Dull et al J Virol. 1998 Nov . 72(11):8463-71. | Addgene #12259 |
| pX330 | pX330-U6-Chimeric_BB-CBh-hSpCas9 | A human codon-optimized SpCas9 and chimeric guide RNA expression plasmid. Used to clone in *VCL* targeting sgRNA for MS2 cassette insertion. | Cong et al Science. 2013 Jan 3. | Addgene #42230 |
| pRPG200 | pLTUB_MCP-HaloTag-GB1 | Lentiviral transfer vector for expressing MCP-HaloTag-GB1 for labeling of MS2-tagged RNAs. | This study. | Available upon request. |

### Supplementary Table 7: Details of transfections performed in this study.

| Transfection | Plate | Plasmid Amounts | Lipofectamine 3000 | P3000 | Opti-MEM |
|---|---|---|---|---|---|
| Generation of U2OS_dSaCas9-ArrayG/N + mwGFP | 12-well plate | 400 ng pRPG001 , 400 ng pRPG102, 200 ng pbTP | 2 uL | 2 uL | 100 uL |
| Generation of 3rd gen lentivirus for sgRNA delivery | 6-well plate | 850 ng pMDLg/pRRE, 850 ng pRSV-Rev, 550 ng pMD2.G, 750 ng 3rd Gen Lentiviral Transfer Vector | 7 uL | 6 uL | 500 uL |
| Transient transfection of sgRNAs for imaging | 8-well ibidi u-Slide | 250 ng sgRNA plasmid | 0.5 uL | 0.5 uL | 12.5 uL |

### Supplementary Table 8: Specifications of PCR for DAP array cloning.

| | Fragment Amplification | Vector Linearization |
|---|---|---|
| Template | pAR104 | pAR103 |
| FWD primer[a] | GTTATAGTACTCTGGAAACAGAATC | GTTATAGTACTCTGGAAACAGAATC |
| REV primer[a,b] | AGGGGGCACCCGGAT | AGGGGGCACCCGGAT |
| Initial denaturation | 98°, 30 sec | 98°, 30 sec |
| Denaturation | 98°, 10 sec | 98°, 10 sec |
| Annealing | 62°, 10 sec | 67°, 20 sec |
| Extension | 72°, 5 sec | 72°, 45 sec |
| Cycles | 35 | 35 |
| Final extension | 72°, 2 min | 72°, 2 min |
| Expected band size | ~220 bp | ~2200 bp |

[a] The sequence of the primer that binds to the template is shown; the full primer also includes ~half of the sgRNA and BsaI cutsites. [b] Sometimes the sgRNA sequence is complementary to the template and the reverse primer must be shortened to decrease the melting temperature.
